# Oxygen availability and hypoxia-independent action of HIF1α controls human trophoblast maturation and function

**DOI:** 10.64898/2025.12.12.693923

**Authors:** Johanna Lattner, Javier Bregante, Michaela Burkon, Ornella Elezaj, Meritxell Huch, Michele Marass, Claudia Gerri

## Abstract

Placenta progenitor cells, also known as trophoblasts, initially specify at very low oxygen (O_2_) concentrations. Across their differentiation path in the uterine microenvironment, they encounter a wide range of O_2_ levels. Despite previous efforts, dissecting how these rapid and dynamic O_2_ levels are sensed by trophoblast stem cells and transduced via hypoxia inducible factors (HIFs) has been challenging and has led to conflicting conclusions. This is at least in part due to the lack of tractable and reliable methods to model human placental development. Here, by recapitulating the dynamic O_2_ levels of the uterine microenvironment, and by genetically ablating *HIF1α* in human trophoblast organoids, we found that O_2_ availability and HIF pathway independently control trophoblast lineage specification, maturation and function. Specifically, low O_2_ levels promote expansion of extravillous trophoblast (EVT) progenitors independently of HIF1α, while HIF1α is necessary for EVT invasion regardless of O_2_ availability. Altogether, our results reveal a dual regulatory framework that disentangles the role of O_2_ from that of HIF1α, offering a revised view of how O_2_ availability regulates early human placental development.

## INTRODUCTION

The placenta mediates communication and exchange of nutrients, waste products, and oxygen (O_2_) between the mother and fetus. Following implantation, placental progenitors derived from the trophectoderm differentiate and give rise to three major trophoblast cell types (**Figure 1A**). Cytotrophoblasts (CTBs) are the proliferative epithelial stem cell pool of the placenta and give rise to two principal lineages. One is the syncytiotrophoblast (SCTs), a multinucleated cell formed through CTB fusion, in which gas and nutrient exchange occur. The second lineage forms in anchoring villi attached to the maternal decidua, where CTBs form a column of proliferative cytotrophoblasts (cCTBs). As cCBT progress distally, they stop proliferation and differentiate into extravillous trophoblasts (EVTs), which invade the uterus and remodel maternal spiral arterioles to establish maternal blood flow to the placenta^1,2^. This differentiation trajectory involves a series of transcriptionally distinct cCTB states that gradually transition into migratory, mature EVTs^3–5^.

**Figure 1.**
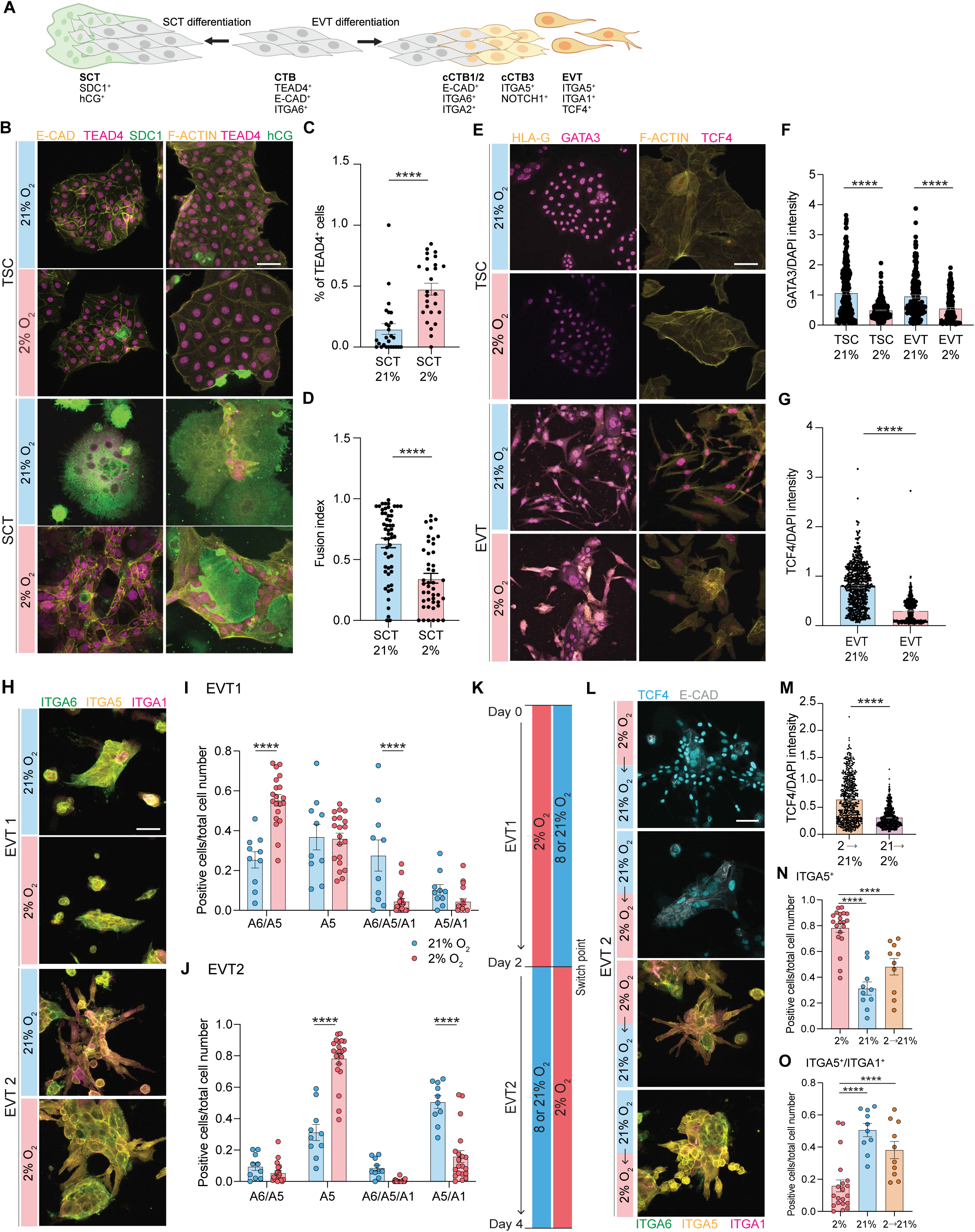
Low O_2_ restricts SCT formation and EVT maturation in 2D cultures. (A) Schematic of cytotrophoblast (CTBs) differentiation into syncytiotrophoblasts (SCTs) and extravillous trophoblasts (EVTs). Created by BioRender. (B) Immunofluorescence images of E-CADHERIN, TEAD4, SDC1, and F-ACTIN, TEAD4, HCG in WT TSCs and SCTs at 21% and 2% O_2_. (C) Quantification of the percentage of nuclear TEAD4-positive cells in SCTs at 21% and 2% O_2_. (*n* = 26 cell colonies for each condition). Statistical analysis by Mann-Whitney U test. (D) Quantification of fusion index in SCTs at 21% and 2% O_2_. (*n* = 53 cell colonies for SCT 21%; *n* = 44 for SCT 2%). Statistical analysis by Mann-Whitney U test. (E) Immunofluorescence analysis of GATA3, HLA-G, and F-ACTIN, TCF4 in WT TSCs and EVTs at 21% and 2% O_2_. (F) Quantification of nuclear GATA3 normalized fluorescence intensity in WT TSCs and EVTs at 21% and 2% O_2_. (*n* = 264 cells for TSC 21%; *n* = 292 for TSC 2%; *n* = 179 for EVT 21%; *n* = 177 for EVT 2%). (G) Quantification of nuclear TCF4 normalized fluorescence intensity in EVTs at 21% and 2% O_2_. (*n* = 607 cells for each condition). Statistical analysis by Mann-Whitney U test. (H) Immunofluorescence analysis of ITGA6, ITGA5 and ITGA1 in WT EVT1 and EVT2 at 21% and 2% O_2_. (I, J) Quantification of cell populations positive for ITGA6, ITGA5, and ITGA1 signal and combination of expression in EVT1 and EVT2 at 21% and 2% O_2_. (*n* = 10 cell colonies for 21%; *n* = 20 for 2%). Statistical analysis by Two-way ANOVA and Tukey’s post-hoc test for multiple comparisons. (K) Schematic of O_2_ switch experiment where differentiating TSCs cultured in EVT1 medium in 2% O_2_ were moved to 8% or 21% O_2_ on day 2 before changing to EVT2 medium, and vice versa. (L) Immunofluorescence analysis of TCF4, E-CADHERIN, and ITGA6, ITGA5, ITGA1 in WT EVT2 in O_2_ switch experiment. (M) Quantification of normalized nuclear TCF4 fluorescence intensity in O_2_ switch experiment. (*n* = 569 cells for each condition). Statistical analysis by Mann-Whitney U test. (N, O) Quantification of ITGA5^+^, and ITGA5^+^/ITGA1^+^ populations in EVT2 from (H) and in O_2_ switch experiment from (L). (*n* = 10 cell colonies for 21% and switch experiment; *n* = 20 for 2%). Statistical analysis by Two-way ANOVA and Tukey’s post-hoc test for multiple comparisons. A5 = ITGA5; A6 = ITGA6; A1 = ITGA1. Error bars indicate mean +/- SEM. **** p<0.0005. Scale bars, 100 μm.

Among other features, the EVT branch is marked by sequential changes in integrin expression signature, which provides a useful indicator of developmental progression. CTBs and proximal cCTBs express mainly integrin α6 (ITGA6), whereas distal cCTBs express integrin α5 (ITGA5) which is maintained in migratory EVTs together with integrin α1 (ITGA1)^6–8^. This transition is often described as a partial epithelial-to-mesenchymal transition (EMT)^9,10^. During this process, CTBs loosen cell-cell contacts, remodel their actin cytoskeleton, and secrete extracellular matrix (ECM)-modifying proteins, enabling acquisition of a migratory EVT phenotype. Key EMT regulators include the transcription factors SNAIL, SLUG, TWIST1, MSX2 that are controlled by the FGF, TGFβ and WNT pathways. Together they drive downregulation of epithelial cell adhesion molecules (e.g., E-CADHERIN and ZO-1), upregulation of cytoskeletal components promoting motility (e.g., TLN1, TNS3, ACTN1, FMN1), and changes in ECM deposition and degradation (e.g., COL1A1, COL1A2, FN1, MMPs) to remodel their environment^5,11–13^. Because EMT and cell migration are energetically demanding, cells must adapt their metabolism during differentiation. In SCTs, differentiation is accompanied by a shift from glycolysis to oxidative phosphorylation (OxPhos)^14,15^, but direct evidence for a metabolic shift during EVT maturation is still missing.

*In vivo* trophoblast differentiation occurs under low O_2_ conditions, approximately at 2% O_2_ during the first trimester and rising to around 8% during the second and third trimesters^16–18^. These low O_2_ levels are crucial for CTB maintenance by regulating metabolic, growth, and differentiation pathway^19–22^. Low O_2_ has also been reported to inhibit SCT formation^23,24^. In humans this poses severe consequences for pregnancy outcomes. One example is the pregnancy disorder intrauterine growth restriction, a pregnancy disorder characterized by decreased syncytialisation^25^. Despite extensive investigation, the influence of low O_2_ on EVT differentiation remains unclear, with studies presenting contradicting results. Some reports indicate that low O_2_ enhances cCTB formation^22,26,27^, whereas others reported the opposite effect^28–30^. Similarly, migratory EVT differentiation has been reported to be both restricted^22,26,27^ and enhanced in low O_2_^23,31^. These conflicting results likely stem, on one hand, from the wide range of experimental models used (from primary villous explant and primary cells to immortalized placenta cancer cell lines) and, on the other hand, from inconsistent terminology and benchmarking used to describe the various stages of EVT differentiation^32^. Recent efforts have begun to reconcile these discrepancies, suggesting that low O_2_ may support cCTB expansion^31,32^. Resolving these discrepancies is crucial to elucidate the role of O_2_ in placenta development and to understand how impaired trophoblast migration and inadequate remodeling of maternal arteries are associated with serious pregnancy disorders, such as preeclampsia or intrauterine growth restriction^33^.

In low O_2_ conditions, cells activate the hypoxia inducible factor (HIF) pathway, by stabilizing the HIFα subunit, which dimerizes with HIFβ subunit upon translocating to the nucleus and binds to the DNA allowing transcriptional activation of downstream targets^34^. Evidence from mouse models indicates that HIFα is required for proper placenta development^35,36^. However, mouse and human placenta development differ substantially in morphology, invasion behavior, and trophoblast lineages^37^. Further, critical molecular pathways in mouse placenta development are not fully conserved in other species, including humans^38–40^. So far, downregulation of HIFβ subunit has revealed that EVT differentiation is supported by HIF activity. However, a clear understanding of the exact role of HIF1α, at which step of EMT progression during EVT differentiation it acts, and how it ties to O_2_ availability is still lacking. Moreover, no direct evidence for a metabolic switch from cCTBs to migratory EVTs exists, and the process of differentiation is not clearly distinguished from EVT function.

Here, we investigate the effect of physiologically low (2%) or atmospheric (21%) O_2_ on differentiation of *in vitro* cultures of *wild-type* and *HIF1α*-deficient (*HIF1α^-/-^*) TSC lines and organoids, which can give rise to all of the trophoblast lineages. By modulating O_2_ availability throughout differentiation in these models, we show the contribution of O_2_ levels to specific steps of both SCT and EVT differentiation. Low O_2_ level prevents both SCT and EVT maturation. Strikingly, this O_2_-dependent inhibition is reversible, as exposing immature EVTs to high O_2_ levels is sufficient to trigger their lineage specification program and generate fully migratory EVTs. Using single cell RNA-sequencing (scRNA-seq) and quantitative immunofluorescence imaging, we dissect the progression of cCTB stages leading up to migratory EVTs. In *wild-type* organoids, low O_2_ expands the pool of cCTBs independently of *HIF1α*. *HIF1α^-/-^*organoids show similar progression of differentiation through populations of cCTB and EVT to the *wild-type*, but *HIF1α^-/-^* EVTs fail to migrate into the surrounding matrix. This hints at a cell type-specific functional defect rather than a complete failure to differentiate. We show that these defects are associated with impaired actin cytoskeleton remodeling and focal adhesion formation. Overall, our work dissects the role of O_2_ and O_2_-dependent and -independent HIF response in human trophoblast lineage differentiation and provides new insights into human placental development in physiology and disease.

## RESULTS

### Low O_2_ levels prevent maturation of syncytiotrophoblasts and extravillous trophoblasts

In our *in vitro* system, we can follow the differentiation of TSCs^41^, representing the CTB pool, into SCTs or EVTs (**Figure 1A**). Although most *in vitro* trophoblast systems are cultured under atmospheric O_2_ (21%), physiological O_2_ levels in the uterus and developing placenta are substantially lower. This discrepancy is particularly relevant for placenta biology, for which rapid shifts in O_2_ tension are a defining feature of early development^16,18^. To mimic the initial uterine environment characterized by low O_2_ levels, we cultured human TSCs and trophoblast organoids at 21% O_2_ and compared them with those grown at 2% O_2_. In 2D culture, TSC colony morphology remained largely comparable between the two conditions; however, the general trophoblast marker GATA binding protein 3 (GATA3) was markedly reduced at 2% O_2_ (**Figure 1E, F**). Under 21% O_2_, TSCs efficiently differentiate into SCTs, forming large multinucleated syncytia accompanied by a reduced E-CADHERIN (E-CAD) signal, changes in F-ACTIN organization and a high fusion index (**Figure 1B-D**). This fate transition is further supported by high expression of human chorionic gonadotrophin (hCG) and SYNDECAN1 (SDC1), well-established SCT markers^42^, and a loss of nuclear TEA domain transcription factor (TEAD4), a stem cell marker^43^ (**Figure 1B, C**). In contrast, at 2%, O_2_, syncytialization is impaired. Colonies retain a low fusion index, strong E-CAD, F-ACTIN and nuclear TEAD4 expression, with only scattered syncytial cells expressing hCG and SDC1 (**Figure 1B-D**).

We next differentiated TSCs into EVTs and found substantial morphological and molecular differences reflecting distinct levels of maturation. At 21% O_2_, cells are elongated, characteristic of fully differentiated EVTs. In contrast, at 2% O_2_, cells appear less elongated with residual stem cell colonies (**Figure 1E**). EVTs generated at 21% O_2_ express HLA-G, a canonical EVT marker, and show high levels of TCF4 ^44^, a WNT transcription factor important for EVT differentiation and invasion^3,44^. At 2% O_2_, TCF4 expression is reduced, while HLA-G is highly expressed (**Figure 1E, G**). Moreover, cells at 2% O2 retain prominent cortical F-ACTIN staining, indicating regions of incomplete EVT differentiation (**Figure 1E**). EVT differentiation proceeds through sequential changes in integrin expression. Integrin expression change as EVTs mature and can be used to characterize maturation state^6^. To characterize how O_2_ levels influence these temporal transitions, we performed a time-course experiment and sampled cells at day 2 (EVT1) and 4 (EVT2) of the differentiation protocol. We also included 8% O_2_ to reflect the physiological O_2_ range encountered as the placenta transitions from the first to second trimester and to model the highest level of O_2_ experienced by EVTs during decidual invasion^16^. At both 21% and 8% O_2_, EVT1 cells express ITGA6 and ITGA5 consistent with a cCTB identity, whereas by EVT2, have reduced ITGA6 expression and increased ITGA1, characteristic of migratory EVTs (**Figure 1H, Figure S1A**). In contrast, at 2% O_2_, cells retain ITGA5 expression but fail to switch efficiently from ITGA6 to ITGA1 (**Figure 1H, Figure S1A**). We confirmed these data by assaying integrin abundance through image-based quantification. At EVT1, cells positive for ITGA5 and ITGA1 (ITGA5^+^/ITGA1^+^) were reduced at 2% O_2_ compared with 8% and 21% O_2_ (**Figure 1I, Figure S1B**). At EVT2, cells cultured in 2% O_2_ display a high proportion of cells expressing only ITGA5 (ITGA5^+^), while cultures at 8% and 21% O_2_ show greater ITGA5^+^/ITGA1^+^ cell number, consistent with an overall more complete EVT maturation (**Figure 1J, Figure S1C**).

Together, these findings reveal a striking effect of low O_2_ levels on trophoblast differentiation: the two major trophoblast lineages fail to achieve the terminal maturation states normally observed at high O_2_ levels.

### Trophoblast organoids exposed to physiological O_2_ levels exhibit reduced syncitia and less invasive EVTs

Trophoblast organoids provide a powerful system to study the dynamics of human placenta development *ex vivo*. Trophoblast organoids grown in Matrigel display an inner core of SCTs with an outer layer of CTBs (called TB-Org), or can be differentiated to form cCTBs and EVTs (called EVT-TB-Org)^45,46^. Thus, they offer the unprecedented opportunity to study the impact of both intrinsic and microenvironmental factors in the formation of the different placental lineages.

To determine the impact of distinct O_2_ tensions on trophoblast differentiation, we first examined SCT marker expression by immunofluorescence. At 21% O_2_, TB-Orgs show an outer layer of TEAD4-positive cells surrounding a large SDC1-positive syncytial core. In contrast, TB-Orgs grown at 2% O_2_ appear smaller and display a reduced SDC1-positive area, as confirmed by quantification (**Figure 2A-C**).

**Figure 2.**
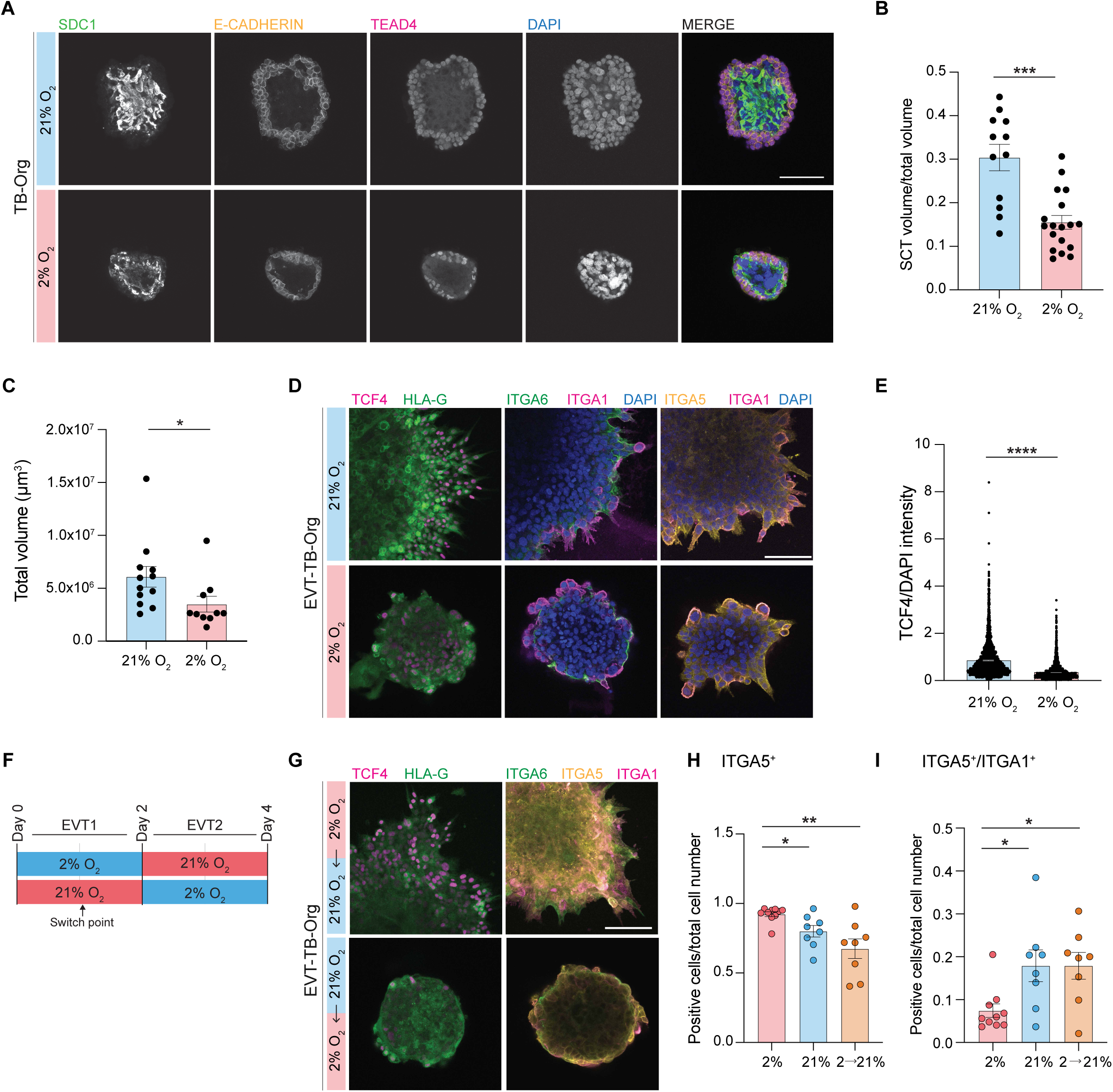
3D trophoblast organoids show loss of proper spatial organization of SCT and EVT progression at 2% O_2_. (A) Immunofluorescence analysis of SDC1, E-CADHERIN, TEAD4, and DAPI signal in WT TB-Orgs at 21% and 2% O_2_. (B) Quantification of SCT volume over total organoid volume in TB-Orgs at 21% and 2% O_2_. (*n* = 12 organoids for 21%; *n* = 18 for 2%). Statistical analysis by Mann-Whitney U test. (C) Quantification of total organoid volume in TB-Orgs at 21% and 2% O_2_. (*n* = 12 organoids for 21%; *n* = 10 for 2%). Statistical analysis by Mann-Whitney U test. (D) Immunofluorescence analysis of TCF4, HLA-G, and ITGA1, ITGA6, DAPI, and ITGA1, ITGA5 and DAPI signal in WT EVT-TB-Orgs at 21% and 2% O_2_. (E) Quantification of nuclear TCF4 normalized fluorescence intensity in EVT-TB-Orgs at 21% and 2% O_2_. (*n* = 2740 cells for 21%; *n* = 2152 for 2%). Statistical analysis by Mann-Whitney U test. (F) Schematic of O_2_ switch experiment where EVT-TB-Orgs cultured in EVT1 medium at 2% O_2_ were moved on day1 to 21% O_2_, and vice versa. (G) Immunofluorescence analysis of TCF4, HLA-G, and ITGA1, ITGA6, ITGA5 in WT EVT-TB-Orgs in O_2_ switch experiment. (H, I) Quantification of ITGA5^+^ only, and ITGA5^+^/ITGA1^+^ populations in EVT2 from (D) and in O_2_ switch experiment from (G). (*n* = 8 organoids for 21% and switch experiment; *n* = 10 for 2%). Error bars indicate mean +/- SEM. Statistical analysis by Two-way ANOVA and Tukey’s post-hoc test for multiple comparisons. * p<0.05; ** p<0.01; *** p<0.005; **** p<0.0005. Scale bars, 100 μm.

Upon EVT differentiation, EVT-TB-Orgs cultured at 21% O_2_ show a spatial progression from HLA-G to TCF4 expression: HLA-G-positive cells are located closer to the inner core, while TCF4 is more expressed at periphery of the organoids (**Figure 2D**). At 2% O_2_, on the contrary, although both TCF4 and HLA-G were present, their spatial organization was disrupted, and EVTs appear less invasive (**Figure 2D, E**). We also analyzed integrin localization in EVT-TB-Orgs. ITGA6 appears similar at 21% and 2% O_2_. ITGA5 expression is higher at 2% O_2_, while ITGA1, which is mostly cytoplasmic and enriched at invasion tips of migratory EVTs at 21% O_2_, is instead detected in the membrane of rounded cells at 2% O_2_, suggestive of incomplete EVT differentiation (**Figure 2D**). We quantified the integrin profiles and EVT-TB-Orgs cultured in 2% O_2_ contained a higher proportion of cells being ITGA5^+^ and fewer ITGA5^+^/ITGA1^+^ cells (**Figure 2H, I**), consistent with our 2D findings.

Overall, organoid architecture at low O_2_ exhibits shallower invasion, with few extruding EVTs. In addition, SCT fusion is reduced.

### Exposure to high O_2_ reverses hypoxia-induced suspension of trophoblast lineage progression

To further model the dynamic levels of O_2_ of the uterine microenvironment encountered by trophoblasts from the beginning of their specification to their differentiation into EVTs and SCTs, we exposed trophoblasts to sequential changes in O_2_ levels. Specifically, we first tested whether 2D trophoblasts cultured initially in 2% O_2_ could resume EVT maturation upon transfer to higher O_2_ concentration (21% or 8%) (**Figure 1K**). We observed that cells experiencing a shift in O_2_ from 2% to 8% or 21% O_2_ upregulate TCF4 and ITGA1 (**Figure 1L-M, Figure S1F-H**). To assess the plasticity of this phenotype and the sensitivity of cells to sudden O_2_ tension reduction, we transferred cells from 21% or 8% O_2_ to 2% O_2_ (**Figure 1K**). Interestingly, cells differentiated in 21% O_2_ show a substantial amount of ITGA5^+^/ITGA1^+^ cell population (**Figure S1D, E**) even after exposure to 2% O_2_, suggesting a resilience in cell commitment. On the other hand, cells grown at 8% O_2_ and transferred to 2% O_2_ show a marked reduction in ITGA5^+^/ITGA1^+^ cells and an expansion of ITGA5^+^ cells, comparable to continuous culture in 2% O_2_ (**Figure S1F-H**). These results indicate that trophoblasts differentiated under intermediate O_2_ conditions retain substantial phenotypic plasticity. For the purpose of our investigation, we focused our attention on the shift between 2% O_2_ and 21% O_2_ in 3D, because EVTs appear more invasive in TB-Org at 21% O_2_ than at 8% O_2_. We reproduced the 2D phenotype in the EVT-TB-Orgs. EVT-TB-Orgs transferred from 2% O_2_ to 21% O_2_ show more ITGA5^+^/ITGA1^+^ cells comparable with those continuously cultured at 21% O_2_ (**Figure 2G-I**). Conversely, organoids shifted from 21% O_2_ to 2% O_2_ exhibited an expanded ITGA5^+^ cell population and reduced ITGA5^+^/ITGA1^+^ population, as expected from organoids cultured in 2% O_2_ throughout (**Figure S1I, J**). Altogether, we find that insufficient O_2_ availability for trophoblasts in 2D or 3D conformation halts their maturation in a transitory state. Importantly, this phenotype can be rescued by providing higher O_2_ levels.

### Single-cell transcriptomics reveals expansion of EVT progenitors under low O_2_ conditions

To gain deeper understanding of the phenotypes observed in our 2D and 3D models and lineage composition under different O_2_ levels, we performed scRNA-seq on TB-Orgs and EVT-TB-Orgs grown at 2% O_2_ and 21% O_2_ (**Figure 3A**). The scRNA-seq analysis reveals eight major clusters: two CTB-like populations (CTB1 and CTB2), three cCTB-like populations (cCTB1, cCTB2, cCTB3), two SCT progenitor-like populations (SCTp1 and SCTp2), and one EVT population (EVT) (**Figures 3B**, **S2A-C**). CTB clusters expressed canonical stem-associated markers, including *TEAD4* and *VGLL1,* and proliferation markers (e.g. *TOP2A*) (**Figure 3D**). The three cCTB clusters reflected a developmental trajectory. The cCTB1 population retain the expression of CTB-associated genes, e.g. *EGFR* and *EPCAM*, while upregulating early cCTB markers, e.g. *TJP1* and *SOX15*. The cCTB2 population represent an intermediate state strongly marked by expression of cCTB markers, such as *ITGA2*, *TAGLN*, *TJP1*, *L1CAM*, and *SOX15*. The cCTB3 cluster show higher expression of *ITGA5*, *FN1*, *LAMA5*, *MMP14*, as well as *ADAM8* and *NOTCH1*. The EVT population, while retaining some cCTB3 transcriptional features, were distinguished by downregulation of *NOTCH1* and robust expression of classical EVT markers, e.g. *MMP2*, *TCF7/L2* (gene name for TCF4), *HLA-G*, *ITGA1*, *NOTUM*, *ADAM12*, *TFGB1*, *NOTCH2* and *SERPINE2* (**Figure 3D**). Although scRNA-seq does not capture multinucleated SCTs, it reliably identifies SCT progenitors (SCTp), expressing *MCM10*, *CYP19A1*, *ERVV-1*, *ERVV-2*, and *SDC1* (**Figure 3D**). Density plots for *ITGA6*, *ITGA5*, and *ITGA1* show the integrin progression from CTBs to EVTs (**Figure 3C**).

**Figure 3.**
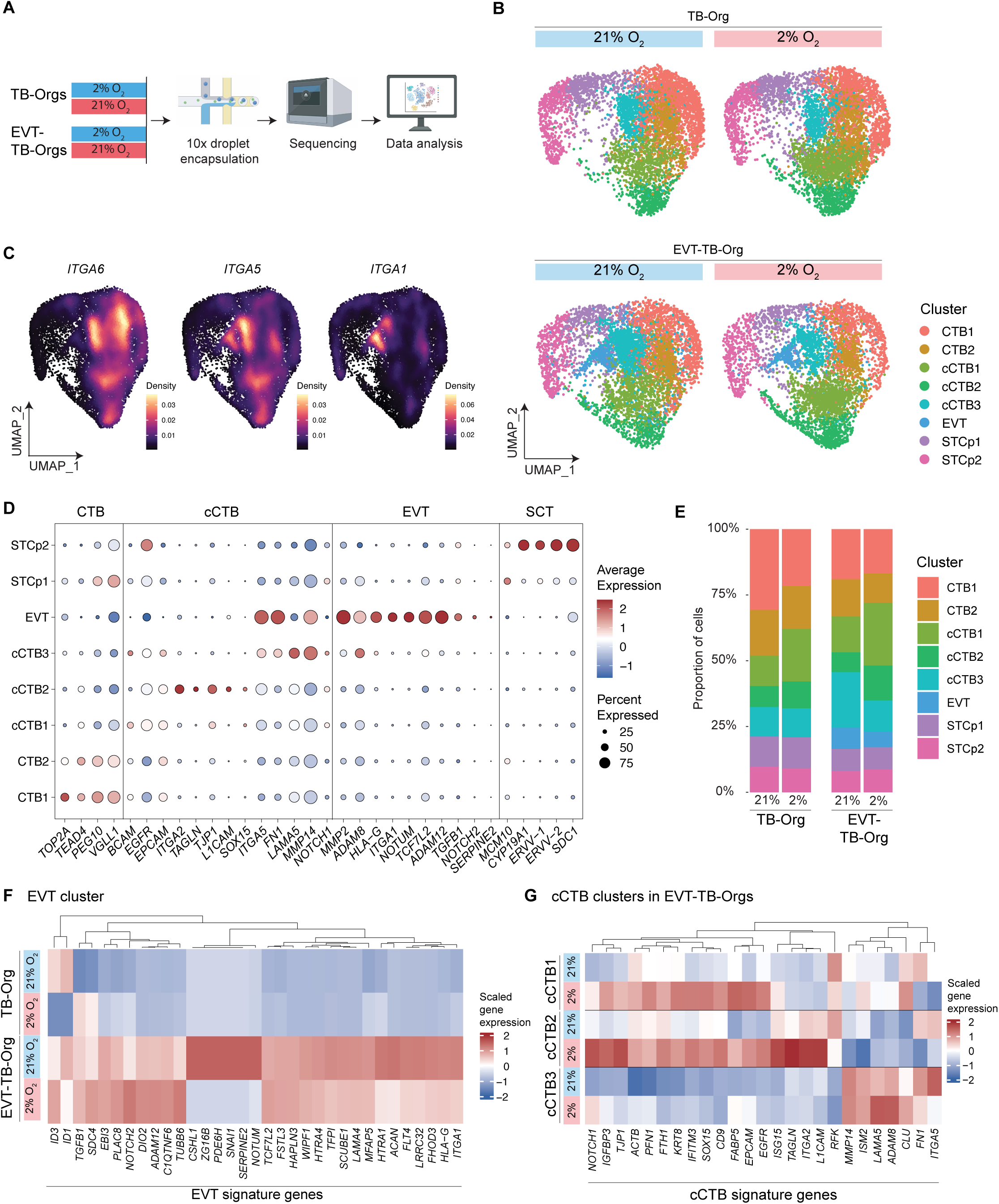
Single cell transcriptomics reveals expansion of cCTB population under low O_2_ conditions. (A) Experimental scheme. WT TB-organoids (TB-Orgs) and EVT-differentiated TB-organoids (EVT-TB-Orgs) were cultured under 21% O₂ and 2% O₂ conditions, dissociated into single cells, and processed for scRNA-seq. (B) Uniform Manifold Approximation and Projection (UMAP) of 23,279 cells, split by condition, showing TB-Orgs and EVT-TB-Orgs at 21% and 2% O₂ and colored by cell type (TB-Orgs 21%: 5,815 cells; TB-Orgs 2%: 6,411 cells; EVT-TB-Orgs 21%: 6,071 cells; EVT-TB-Orgs 2%: 4,982 cells). (C) UMAP feature-density plots dynamic expression patterns of key integrins involved in the transition from CTBs to EVTs in all four datasets. (D) Dot-plot showing gene expression of representative signature markers for each trophoblast lineage. The dot size corresponds to the ratio of cells expressing the gene in the population. The colour scale corresponds to the cell type averaged gene expression level. (E) Cell proportions of clusters identified in TB-Orgs and EVT-TB-Orgs at 21% and 2% O_2._ (F) Heatmap showing scaled gene expression of EVT signature genes in the EVT cluster in TB-Orgs and EVT-TB-Orgs at 21% and 2% O_2._ (G) Heatmap showing scaled gene expression of cCTB signature genes in the cCTB clusters (cCTB1, cCTB2 and cCTB3) in EVT-TB-Orgs at 21% and 2% O_2._

We next profiled the cell proportion changes across the four conditions. Interestingly, across both TB-Orgs and EVT-TB-Orgs we observed an expansion of cCTB1 and cCTB2 populations under 2% O_2_. In TB-Orgs, this expansion occurred at the expense of CTBs, whereas in EVT-TB-Orgs, it resulted in only a minor reduction of CTBs but led to a more pronounced depletion of cCTB3, EVTs and SCT progenitors (**Figure 3E**). EVT signature gene expression was reduced in 2% O_2_, including *NOTUM* (**Figure 3F**). Genes involved in EMT, such as *SNAI1* and *SERPINE2*^47,48^, were also downregulated (**Figure 3F**). Analysis of cCTB signature across the three cCTB populations shows a clear upregulation of cCTB markers in cCTB1 and cCTB2 at 2% O_2_, whereas cCTB3 did not show a comparable increase (**Figure 3G**). Similarly, SCT progenitor clusters exhibited reduced expression of SCT identity genes, such as *ERVV-1*, *ERVV-2* and *INSL4*, under 2% O_2_ (**Figure S2D**). Overall, these transcriptomic data show that low O_2_ levels promote expansion of cCTBs, while suppressing progression toward fully differentiated EVT and SCT lineages. This shift is accompanied by reduced expression of EMT- and invasion-associated genes that are essential for EVT function.

### Hypoxia signaling is activated during EVT specification even under high O_2_ levels

To examine how O_2_ levels influence hypoxia-associated pathways during trophoblast differentiation, we assessed hypoxia-related gene signatures across our scRNA-seq dataset. PROGENy analysis^49^ reveals detectable hypoxia pathway activity following EVT specification even at 21% O_2_, in particular within cCTB1 and cCTB2 (**Figure 4A**). We then looked for the expression of HIF1α targets and observed that the HIF1α regulon was upregulated in cCTB1 and cCTB2 in EVT-TB-Orgs at 21% O_2_ compared to 2% O_2_ (**Figure 4B**). Interestingly, HIF1α protein is detectable in 2D TSCs even at 21% O_2_ both with immunofluorescence and western blot analysis (**Figure S3C, D**). In EVT-TB-Orgs, HIF1α was localized in the nuclei of trophoblast cells adjacent to extruding EVTs in 21% O_2_, and accumulated primarily in the outer layer of organoids grown in 2% O_2_ (**Figure 4C**).

**Figure 4.**
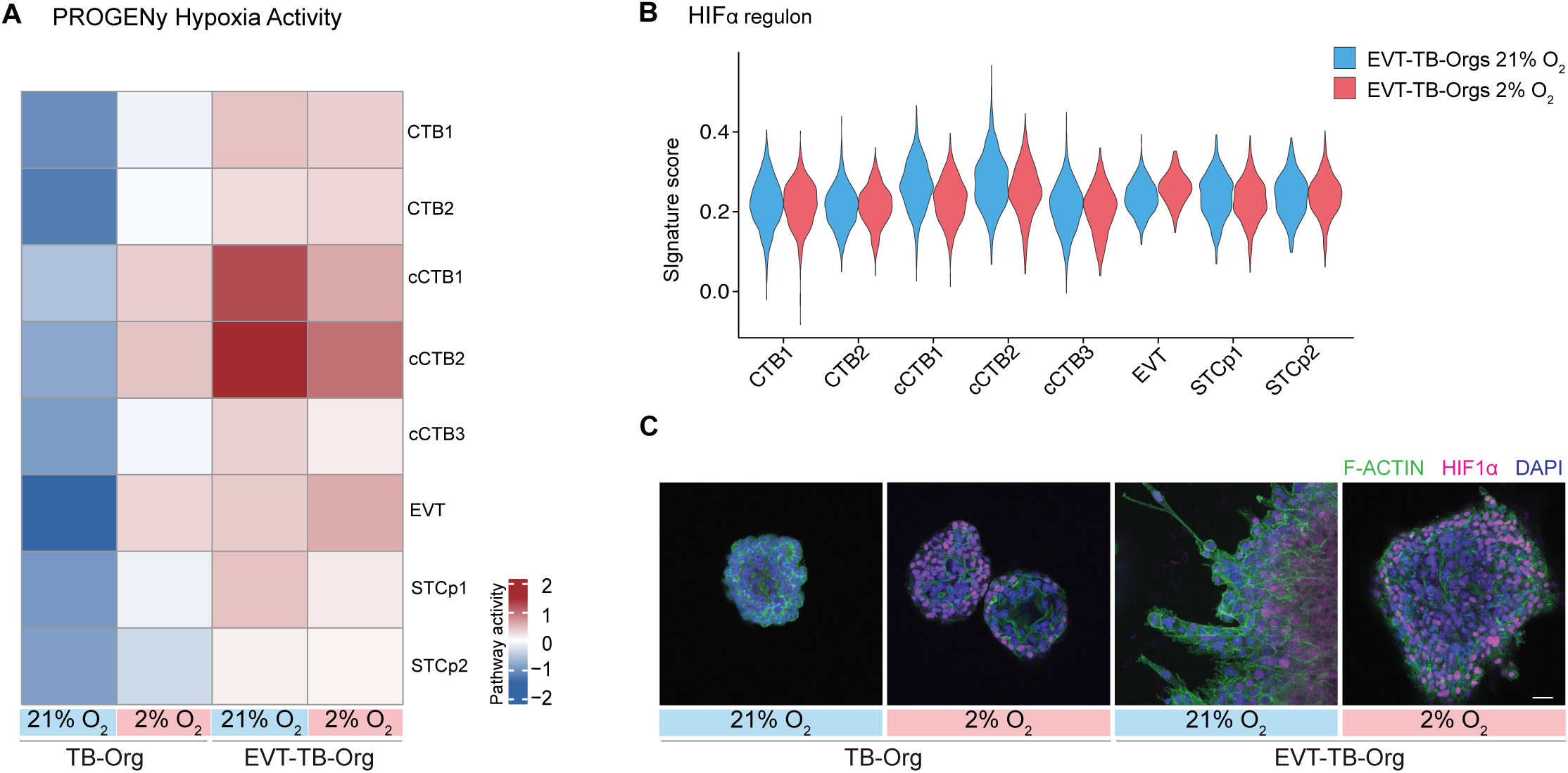
HIF1α is active at 21% O_2_ in EVT-TB-Orgs. (A) PROGENy-based hypoxia pathway across the different clusters in WT TB-Orgs and EVT-TB-Orgs at 21% and 2% O_2_. (B) Violin plot showing HIF1α regulon activity for each cell type in WT EVT-TB-Orgs at 21% and 2% O_2_. (C) Immunofluorescence analysis of HIF1α, F-ACTIN and DAPI signal in WT TB-Orgs and EVT-TB-Orgs at 21% and 2% O_2_. Scale bar, 100 μm.

These data suggest that HIF1α may have a role in EVT differentiation, maturation, or function at low O_2_ levels, but its activation at 21% O_2_ indicates a potential role in EVT differentiation independently of O_2_ levels.

### HIF1α knockout impairs EVT function even at high O_2_ levels

To investigate the specific function of HIF1α and determine whether the expansion of cCTBs observed in low O_2_ depends on this master hypoxia regulator, we generated mutant trophoblast cell lines. Using CRISPR-Cas9-mediated genome editing, we generated *HIF1α* mutant (*HIF1α^-/-^*) CT27 and CT29 TSC lines^41^, that introduce a premature stop codon within bHLH domain (**Figure S3A, B**). By immunofluorescence HIF1α protein signal is lowly expressed in TSCs cultured in 21% O_2_ and strongly induced when they are cultured in 2% O_2_. In contrast, in both CT27 and CT29 *HIF1α^-/-^* cell lines HIF1α is not detected (**Figure S3C**). In addition, we validated our lines by Western Blot, where we observed a strong induction of HIF1α in WT at 2% O_2_ but not in *HIF1α^-/-^* (**Figure S3D**), together confirming HIF1α loss in our mutant cells.

We next analyzed how *HIF1α* loss affects trophoblast lineage specification. *HIF1α^-/-^*TSCs show no major differences compared to WT at both 21% and 2% O_2_, displaying GATA3 reduction at 2% O_2_ (**Figure S4A, B**), as seen in WT (**Figure 1E, F**). Consistent with WT cells (**Figure 1B**), *HIF1α^-/-^* TSCs differentiated into SCTs in 21% O_2_, form multinucleated cells as evidenced by reduced E-CAD intensity and induction of SDC1 expression (**Figure S4C**), which was confirmed by fusion index quantification (**Figure S4D**). Under 2% O_2_, *HIF1α^-/-^* SCTs showed partial induction of SDC1 but failed to form large and extensive syncytia, as reflected by persistent E-CAD staining and reduced fusion index (**Figure S4C, D**). Notably, the fusion defect is more pronounced in *HIF1α^-/-^* then WT (**Figure S4D**). TB-Orgs appear similar between WT and *HIF1α^-/-^* based on cellular organization and presence of TEAD4 and SDC1, where both show a reduction of SCT volume and overall size at 2% O_2_ (**Figure S4E-G**).

We next differentiated *HIF1α^-/-^* TSCs into EVTs. Compared to WT, *HIF1α^-/-^* cells fail to acquire the characteristic elongated morphology under both 21% and 2% O_2_ and display reduced levels of TCF4 and HLA-G (**Figure 5A-C**). Similarly to the WT, *HIF1α^-/^*^-^ cultures show an expansion of ITGA5^+^ cells and a reduction of ITGA5^+^/ITGA1^+^ cells at 2% relative to 21% O_2_ (**Figure D-F**). Notably, compared to WT, *HIF1α^-/-^* colonies contain a higher amount of ITGA5^+^ cells in both 21% and 2% O_2_, and the ITGA5^+^/ITGA1^+^ cell population is reduced (**Figure 5D-F**). Consistent with the 2D observations, *HIF1α^-/-^* EVT-TB-Orgs do not display migratory EVTs in either O_2_ conditions, as shown by brightfield image (**Figure 5G**). This phenotype is accompanied by marked reduction in HLA-G expression and only a minimal TCF4 expression (**Figure 5G, J**). As shown before (**Figure 2D**), WT EVT-TB-Orgs, particularly in 21% O_2_, show a clear progression from HLA-G-positive cells closer to the organoid core towards TCF4-positive cells at the surface, a pattern absent in the mutants (**Figure 5G**). In addition, TCF4 is reduced in both WT and *HIF1α^-/-^* EVT-TB-Orgs at 2% O_2_, with overall lower levels in the mutants (**Figure 5J**). Similarly, both WT and *HIF1α^-/-^* EVT-TB-Orgs at 2% O_2_ show expanded ITGA5^+^ cell population and reduced ITGA5^+^/ITGA1^+^ cell population (**Figure 5G-I**).

**Figure 5.**
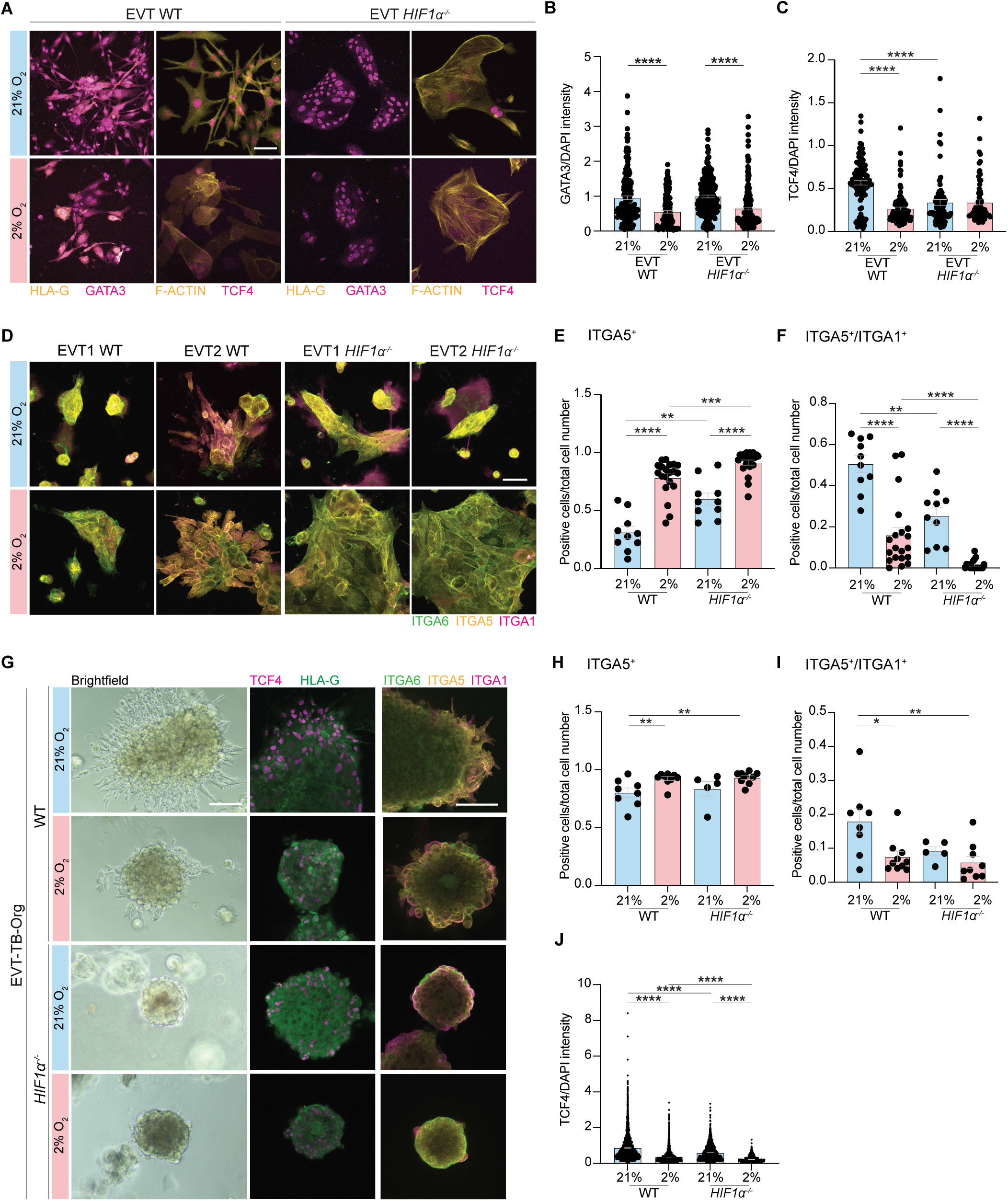
HIF1α has a hypoxia-independent role in EVT lineage progression. (A) Immunofluorescence analysis of GATA3, HLA-G, and F-ACTIN, TCF4 in WT and *HIF1α^-/-^* EVTs at 21% and 2% O_2_. B) Quantification of nuclear GATA3 normalized fluorescence intensity in WT and *HIF1α^-/-^* EVTs at 21% and 2% O_2_ (WT data also shown in Figure 1). (*n* = 180 cells for WT 21%; *n* = 177 for WT 2%; *n* = 191 for *HIF1α^-/-^* 21%; *n* = 207 for *HIF1α^-/-^* 2%). (C) Quantification of nuclear TCF4 normalized fluorescence intensity in WT and *HIF1α^-/-^* EVTs at 21% and 2% O_2_ (WT data also shown in Figure 1). (*n* = 105 cells for WT 21%; *n* = 108 for WT 2%; *n* = 74 for *HIF1α^-/-^*21%; *n* = 72 for *HIF1α^-/-^* 2%). (D) Immunofluorescence analysis of ITGA6, ITGA5, ITGA1 in WT and *HIF1α^-/-^* at EVT1 and EVT2 at 21% and 2% O_2_. (E, F) Quantification of ITGA5^+^ only, and ITGA5^+^/ITGA1^+^ populations in WT and *HIF1α^-/-^* in EVTs at EVT2 at 21% and 2% O_2_ (WT data also shown in Figure 1). (*n* = 10 cell colonies for WT 21%; *n* = 20 for WT 2%; *n* = 10 for *HIF1α^-/-^* 21%; *n* = 20 for *HIF1α^-/-^* 2%). (G) Brightfield images and immunofluorescence analysis of TCF4, HLA-G, and ITGA6, ITGA5, ITGA1 in WT and *HIF1α^-/-^* EVT-TB-Orgs at 21% and 2% O_2_. (H, I) Quantification of ITGA5^+^ only, and ITGA5^+^/ITGA1^+^ populations in WT and *HIF1α^-/-^* in EVT-TB-Orgs at 21% and 2% O_2_ (WT data also shown in Figure 2). (*n* = 8 organoids for WT 21%; *n* = 10 for WT 2%; *n* = 5 for *HIF1α^-/-^* 21%; *n* = 10 for *HIF1α^-/-^* 2%). (J) Quantification of nuclear TCF4 normalized fluorescence intensity in WT and *HIF1α^-/-^* EVT-TB-Orgs at 21% and 2% O_2_ (WT data also shown in Figure 2). (*n* = 2740 cells for WT 21%; *n* = 2152 for WT 2%; *n* = 2350 for *HIF1α^-/-^* 21%; *n* = 1124 for *HIF1α^-/-^* 2%). Error bars indicate mean +/- SEM. Statistical analysis by Two-way ANOVA and Tukey’s post-hoc test for multiple comparisons. * p<0.05; ** p<0.01; *** p<0.005; **** p<0.0005. Scale bars, 100 μm.

Collectively, these data demonstrate that even under high O_2_ conditions, *HIF1α^-/-^*organoids exhibit a pronounced defect in EVT function. Despite expressing some mature EVT markers, the mutant displays impaired invasion and altered organoid organization. Notably, while *HIF1α^-/-^* organoids appear morphologically similar to WT organoids cultured in 2% O_2_, both showing reduced protruding EVTs, the underlying molecular mechanisms are likely distinct.

### scRNA-seq of *HIF1α^-/-^*organoids reveals a complex relationship between O_2_ levels and hypoxia response

To better understand the phenotype of *HIF1α^-/-^*, we performed scRNA-seq of *HIF1α^-/-^*TB-Orgs and EVT-TB-Orgs at 21% O_2_ and 2% O_2_. Comparison across all eight datasets revealed the same eight cell clusters previously identified with WT datasets only (**Figure S5**). Remarkably, in both TB-Orgs and EVT-TB-Orgs the expansion of cCTB1 and cCTB2 populations induced by low O_2_ still occurs even in the absence of HIF1α, indicating that this phenotype is low O_2_-dependent but HIF1α-independent. However, at low O_2_ levels, the number of mature EVT was substantially lower in the mutant compared to WT (**Figure 6A**), consistent with our phenotypic observations of round organoids with a shallow surface and reduced EVT marker expression (**Figure 5G-J**). This striking result, corroborated by transcriptomic analysis, indicates that at low O_2_ and without HIF1α EVT differentiation fails. To understand this, we moved our attention to the cell population distribution of *HIF1α^-/-^* organoids at 21% O_2_, expecting a reduced number of EVTs, in light of our previous phenotypic analysis. Interestingly, the number of EVTs in *HIF1α^-/-^* organoids, according to the cluster analysis and assignment to cell lineages, is comparable to WT organoids, despite a reduction in cCTB3 population (**Figure 6A**). We investigate this specific point in the paragraph below. Of note, we also detected an expansion of SCT progenitors in *HIF1α^-/-^* TB-Orgs at 21% O_2_ relative to WT, as partially confirmed by increase in SCT markers in the scRNA-seq (**Figure S4H**), indicating a possible role of *HIF1α^-/-^* in restricting this cell population. Overall, scRNAseq analysis revealed that cell population distribution was largely similar between mutant and WT organoids and that both responded similarly to low O_2_ levels. Importantly, we observed a notable exception of a dramatic loss of mature EVTs in mutant organoids grown at 2% O_2_ compared with WT organoids grown in the same condition.

**Figure 6.**
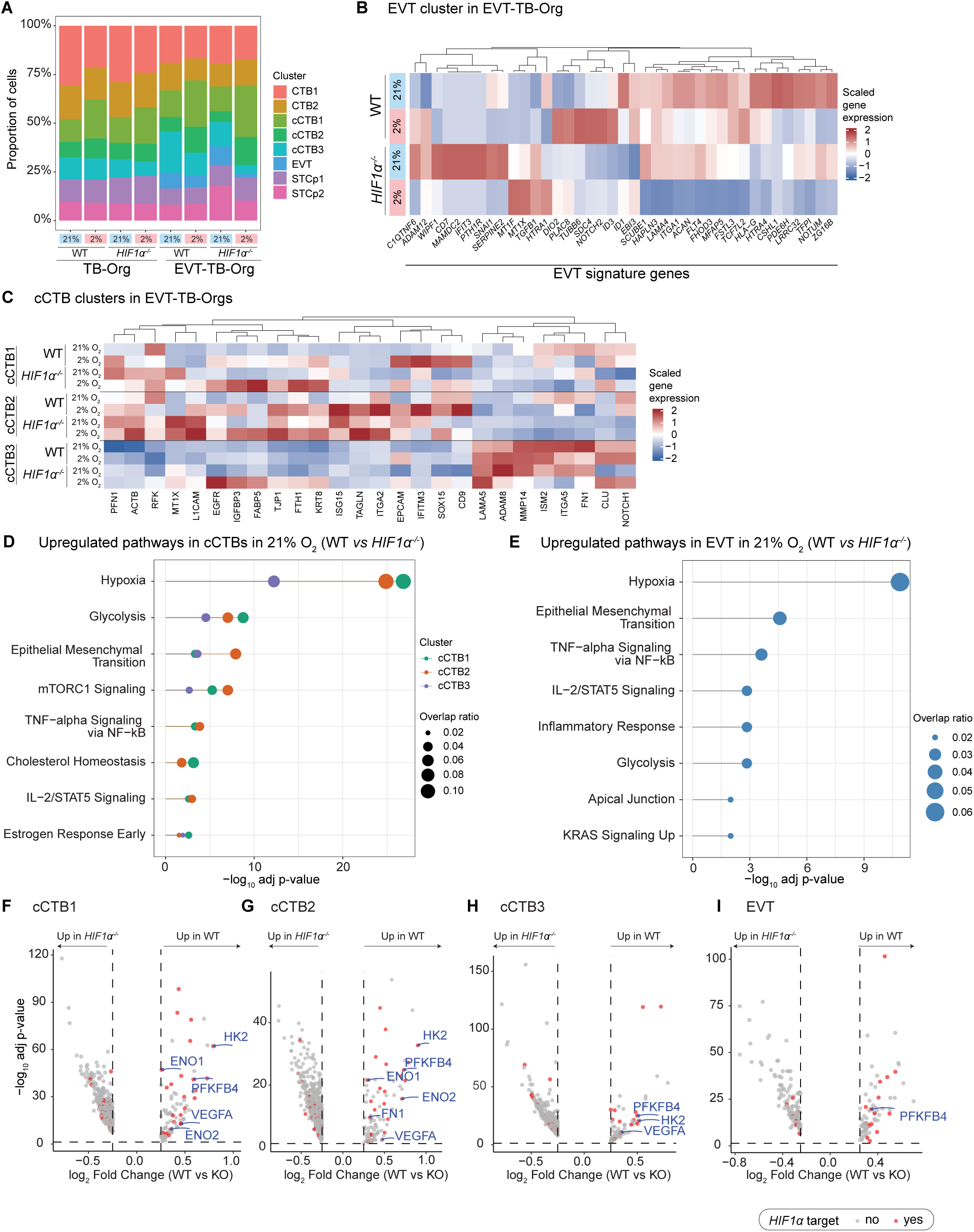
Expansion of cCTBs in low O_2_ is independent of HIF1α. (A) Cell proportions of clusters identified in WT and *HIF1α^-/-^* TB-Orgs and EVT-TB-Orgs at 21% and 2% O_2_. (B) Heatmap showing scaled gene expression of EVT signature genes in the EVT cluster in WT and *HIF1α^-/-^* EVT-TB-Orgs at 21% and 2% O_2_. (C) Heatmap showing scaled gene expression of cCTB signature genes in the cCTB clusters (cCTB1, cCTB2 and cCTB3) in WT and *HIF1α^-/-^* EVT-TB-Orgs at 21% and 2% O_2_. (D, E) Lollipop plot showing the top 8 enriched MSigDB Hallmark pathways for genes upregulated in WT compared to *HIF1α^-/-^* in cCTB populations (cCTB1, cCTB2 and cCTB3) and EVT from EVT-TB-Orgs cultured under 21% O₂. The point size indicates the proportion of overlapping genes in each pathway (overlap ratio). (F-I) Volcano plot showing differentially expressed genes (DEGs) in cCTBs and EVTs in EVT-TB-Orgs cultured at 21% O₂ in WT versus *HIF1α^-/-^*. (|log₂FC| > 0.25, adjusted p < 0.05). Genes annotated as HIF1α targets are highlighted in red.

### HIF1α promotes EMT and focal adhesion formation in EVTs

To reconcile our morphological analysis, that highlighted a lack of invasive EVT in *HIF1α^-/-^* EVT-TB-Orgs at 21% O_2_, with our scRNA-seq analysis, showing a stable number of mature EVT, we focused on analyzing the transcriptional differences of cCTBs and EVTs populations. At first, we look at the EVT signature within the EVT cluster of EVT-TB-Orgs at 21% and 2% O_2_ (**Figure 6B**). *HIF1α^-/-^* EVT-TB-Orgs cultured in 2% O_2_ show a strong decrease in expression of EVT markers. Interestingly, *HIF1α^-/-^* EVT-TB-Orgs cultured in 21% O_2_ retain expression of several canonical EVT markers, such as *NOTUM*, *ITGA1*, *SERPINE2*, *SNAI1*, similar to WT EVT-TB-Orgs cultured at 21% O_2_, but also display some notable differences. Specifically, *HIF1α^-/-^*EVT-TB-Orgs cultured in 21% O_2_ show downregulated expression of *TCF7/L2, HLA-G*, *ID1*, *HTRA4*^50–52^, suggesting that the EVT identity may not be fully preserved in the mutant context (**Figure 6B**). These data suggest that EVTs differentiate and mature even without HIF1α, but might be in a dysfunctional state that results in defective invasion. Strikingly, *HIF1α^-/-^* compared to WT EVT-TB-Orgs cultured in 21% O_2_ exhibit upregulation of *PTHR1*, *MAMDC2*, *CD7*, *IFIT3*, *WIPF3* (**Figure 6B**), the roles of which remains unclear in placenta biology. Analysis of cCTB signature in both WT and *HIF1α^-/-^* EVT-TB-Orgs confirmed the expansion of cCTB1 and cCTB2 clusters at 2% O_2_ (**Figure 6C**). However, cCTB3 population of *HIF1α^-/-^* EVT-TB-Orgs cultured at 2% O_2_ show an increased expression of CTB marker *EPCAM*, and cCTB1 and cCTB2 markers *TJP1* and *EGFR*, suggesting that these cells may be arrested in a more premature state under low O_2_ conditions (**Figure 6C**). Further analysis of EVT and cCTB populations revealed that WT cCTBs and EVTs at 21% O_2_ were enriched for hypoxia-, EMT- and glycolysis-related pathways, compared to *HIF1α^-/-^* EVT-TB-Orgs (**Figure 6D, E**). Differential gene expression (DEG) analysis of HIF1α target genes across cCTBs and EVTs highlight downregulation of key regulators of glycolysis, including hexokinase2 (*HK2*) and 6-phosphofructo-2-kinase-2,6-biphosphatase 4 (*PFKFB4*), as well as EMT/cell migration, such as fibronectin1 (*FN1*)^53–55^ (**Figure 6F-I**).

We then analyzed the expression of key functional EMT markers reflecting different stages of maturations, namely EMT priming, EMT transition and migration across cCTBs and EVTs. This analysis confirmed that the few EVTs observed in mutant organoids at 2% O_2_ are largely defective for EMT transition and migration markers (**Figure 7A**). However, *HIF1α^-/-^* EVT-TB-Orgs at 21% O_2_ showed similar expression of most markers compared with WT, except for notable downregulation of *FN1*, *FMN1* and *FNBP4*, which are involved in actin filament formation and in EMT transition^54,56,57^ (**Figure 7A**). We then examined F-ACTIN, a main component of the cytoskeleton and TALIN1 expression, a key mechanical linker between integrins and actin in focal adhesions^58^, across conditions. In WT EVT-TB-Orgs at 21% O_2_, F-ACTIN expression is well organized into stress fibers, and TALIN1 localizes to protrusive EVTs extending into the matrix. At 2% O_2_, F-ACTIN stress fibers remain visible, but TALIN1 expression is reduced. In *HIF1α^-/-^* EVT-TB-Orgs, these defects are more pronounced, especially at 2% O_2_, where the F-ACTIN stress fibers are lost and accumulate in the cytocortex (**Figure 7B**).

**Figure 7.**
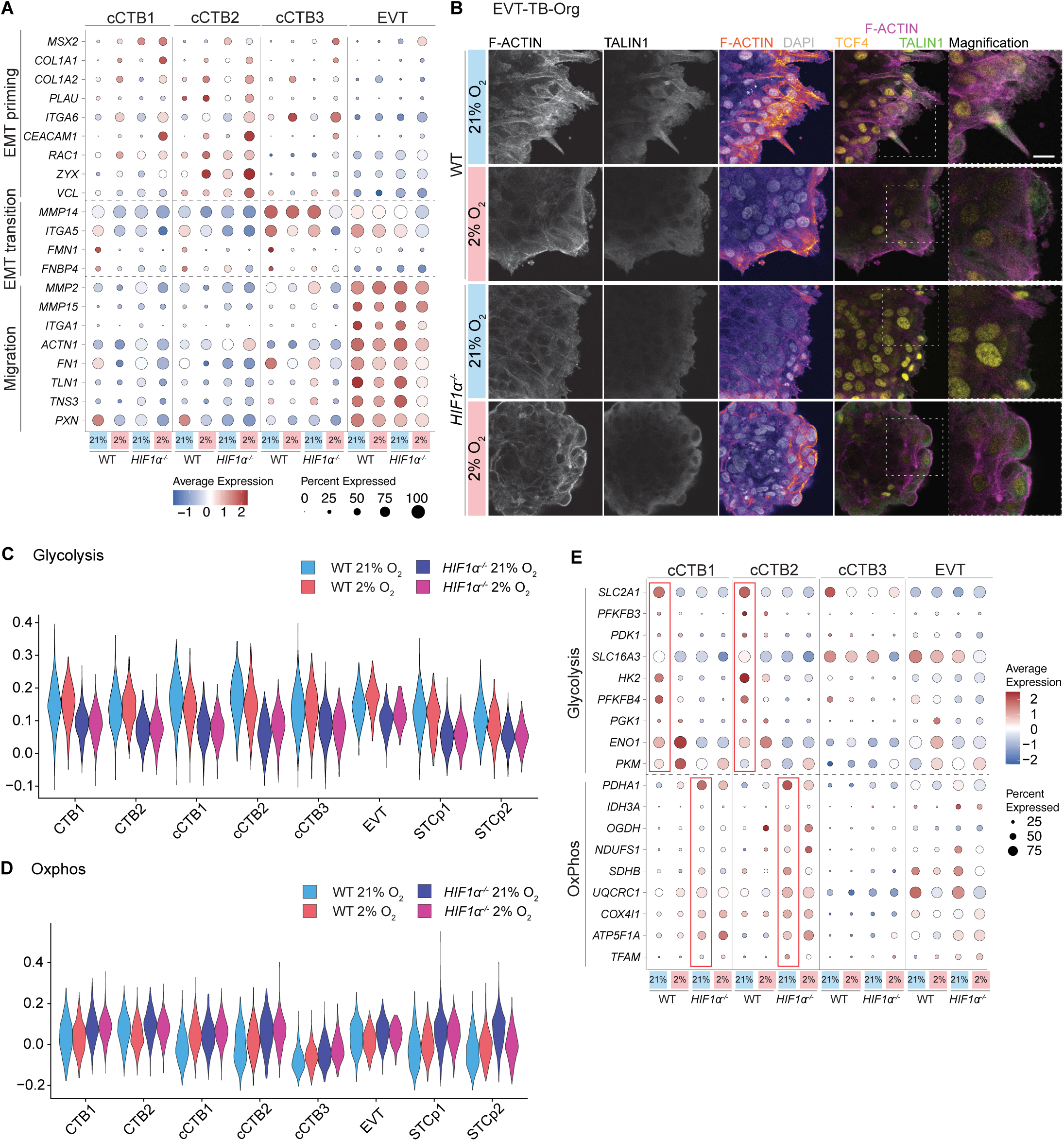
Loss of HIF1α impairs EVT migration by disrupting focal adhesion formation. (A) Dot plot showing scaled gene expression of key EMT and migration-related markers in cCTB1, cCTB2, cCTB3, and EVT in WT and *HIF1α^-/-^* EVT-TB-Orgs at 21% and 2% O_2_. (B) Immunofluorescence of F-ACTIN, TALIN1, DAPI signal in WT and *HIF1α^-/-^* EVT-TB-Orgs at 21% and 2% O_2_. (C, D) Violin plots of glycolysis and OxPhos (oxidative phosphorylation) gene signatures per cell type in WT and *HIF1α^-/-^* EVT-TB-Orgs at 21% and 2% O_2_. (E) Dot plot showing scaled gene expression of key glycolysis and OxPhos markers in cCTB1, cCTB2, cCTB3, and EVT in WT and *HIF1α^-/-^* EVT-TB-Orgs at 21% and 2% O_2_. The dot size corresponds to the ratio of cells expressing the gene in the population. The color scale corresponds to the cell type averaged gene expression level. Scale bar, 100 μm.

DEG-based analysis indicate increased glycolysis activity in WT EVT-TB-Orgs at 21% O_2_, whereas *HIF1α^-/-^* EVT-TB-Orgs show signatures consistent with an energy deficit (**Figure 6E**). Further analysis of glycolysis and oxidative phosphorylation (OxPhos) gene signatures show that all major cell populations, particularly CTBs, cCTBs and EVTs, heavily rely on glycolysis, while *HIF1α^-/-^* cells shift to OxPhos (**Figure 7C, D**). This is further confirmed by examination of specific glycolysis- and OxPhos-related genes (**Figure 7E**). In WT EVT-TB-Orgs at 21% O_2_, cCTB1 and cCTB2 show high expression of glycolytic markers, such as *HK2*, *PFKFB4*, and *SLC2A1* (**Figure 7E**). In contrast, *HIF1α^-/-^* EVT-TB-Orgs at 21% O_2_ display reduced glycolytic gene expression, and increased expression of OxPhos-associated genes, including *PDHA1* (**Figure 7E**).

This energy imbalance could cause defects in energy-intensive processes, including focal adhesion formation, EMT and EVT migration. However, a deeper mechanistic investigation is required in the future to fully elucidate how HIF1α coordinates metabolism and EVT mechanics under high O_2_ levels.

## DISCUSSION

Understanding how trophoblasts, the placenta progenitors, sense and respond to the rapidly changing O_2_ landscape of early pregnancy is central to human development and reproduction. Although the uterine microenvironment is known to be characterized by low O_2_, its precise influence on trophoblast lineage specification, maturation, and function is unclear. This gap in understanding stems in part from reliance on non-physiological *in vitro* models and *in vivo* models that do not fully recapitulate the human placental biology. This is a fundamental question in stem cell and developmental biology, as placenta progenitors represent the first embryonically derived cells to interface with an environment shaped not by the embryo itself, but by a genetically distinct individual, the mother. Here, by modeling the physiological dynamics of O_2_ levels and by generating genetic *HIF1α^-/-^* human trophoblast organoids, we found that O_2_ availability and HIF1α activity exert distinct and non-overlapping roles in guiding trophoblast lineage progression, maturation, and function. First, we identified that exposure of 2D trophoblasts and 3D organoids to low, physiological O_2_ levels drives an expansion of cCTBs, reflecting a transitory cell state during lineage progression. In contrast, standard culturing protocols at atmospheric O_2_ levels drive rapid cell differentiation bypassing the step-wise transitions that normally occur *in vivo*, and therefore obscuring critical intermediate states of lineage progression. Therefore, our findings not only indicate low O_2_ levels act as a bottleneck during trophoblast specification, but highlight the need of culturing cells and organoids in physiological conditions in order to investigate and examine in detail intermediate cell populations and developmental checkpoints that may not be evident at high O_2_ levels. In addition, these immature trophoblast cells retain a notable degree of plasticity, as their transient arrest can be resolved by shifting them to higher O_2_ levels, enabling progression toward full maturation. The ability to rescue this phenotype has potential clinical relevance. In disorders such as preeclampsia, EVTs fail to properly mature and invade the uterine layer, possibly due to prolonged hypoxia^59–61^. Thus, our organoid model could serve as a drug development platform to identify strategies able to unlock EVT maturation, accelerate vascular remodelling in the uterus and improve trophoblast oxygenation through a positive feedback loop. We speculate that compounds targeting metabolic pathways active under high O_2_ contexts, molecules able to restore the energetic imbalance of ATP-deprived cells, or modulators of cellular O_2_ sensing mechanisms beyond HIF1α, could represent promising candidates to promote EVT maturation in low O_2_ conditions.

Unexpectedly, we also found that hypoxia-driven expansion of cCTBs occurs independently of HIF1α. This uncoupling of hypoxia and HIF activity contrasts with the most obvious assumptions and suggests that EVT progenitors expand via other O_2_-sensing mechanisms, possibly through ROS, metabolic rewiring or other HIFs, such as HIF2α/EPAS1 and HIF3α. On the other hand, HIF1α proved to be necessary for EVT function, governing processes such as EMT and invasion even under high O_2_. In fact, *HIF1α^-/-^* EVTs exhibit round morphology, loss of focal adhesions, and reduced HLA-G expression, ultimately failing to invade the matrix despite being exposed to high O_2_ levels. Together, we identified phenotypes that are HIF1α-dependent but hypoxia-independent, as well as phenotypes that are hypoxia-dependent but HIF1α-independent. Thus, rather than acting solely as an O_2_ sensor, HIF1α likely serves as a broader regulatory hub required to meet the energetic, cytoskeletal, and transcriptional demands of EVT function. This disentanglement between HIF1α and hypoxia uncovers a previously unappreciated non-trivial regulatory role of O_2_ availability and HIF1α in trophoblast lineage specification and EVT development.

By integrating phenotypic and transcriptomic data, we propose that HIF1α drives EVT function, rather than their differentiation program per se. The reduction in glycolytic gene expression in *HIF1α^-/-^* EVTs, and the concomitant dysregulation of EMT genes, such as *FMN1* and *FN1*, suggests that HIF1α might be responsible to supply EVTs with sufficient metabolic support for ATP-intensive processes, including EMT, focal adhesion assembly, migration. An alternative, non-exclusive mechanism may involve reactive O_2_ species (ROS), as repression of HIF1α has been previously associated with increased mitochondrial ROS in other contexts, and excessive ROS, accompanied by oxidative stress^62–64^. Moreover, HIF1α has been shown to regulate several genes involved in cytoskeletal structure and EMT, including *FN1*^65^, although whether these represent direct HIF1α targets in trophoblast organoids remains unknown. Investigating in detail the mechanisms linking HIF1α loss and impaired focal adhesion formation in EVTs through energy imbalance, ROS accumulation or dysregulation of direct HIF1α targets warrants future investigation.

One of the most counter-intuitive observations is the apparent synergy between low O_2_ and HIF1α loss. EVTs are in fact substantially reduced in *HIF1α^-/-^* organoids under low O_2_ conditions, a phenotype more severe than either the immaturity caused by hypoxia alone or the dysfunction seen in *HIF1α^-/-^* in high O_2_ levels. Rather than compensating for or reversing hypoxic effects, HIF1α loss appears to exacerbates them. This phenotype likely reflects an adaptive requirement for HIF1α activation during prolonged hypoxia. We hypothesize that *in vivo* EVTs might rely on HIF1α as a survival mechanism under low O_2_ environments, until they reach more oxygenated maternal tissues. If HIF1α function is impaired and/or hypoxia is maintained, EVTs might fail to adapt to low O_2_, resulting in a low number of mature and invasive EVTs under such conditions.

Overall, our findings go beyond the intuitive view of hypoxia as a unidirectional driver of EVT development through HIF1α activation, and instead reveal a complex relationship between hypoxia and HIFs, with physiological low O_2_ being fundamental for proper cCTBs expansion, but with HIF1α being involved in mature EVT function at higher O_2_ levels. More broadly, our results underscore the critical importance of recapitulating physiologically accurate microenvironments for *in vitro* modeling of human development and disease.

### Limitations of the study

The study models O_2_ availability using defined low and high O_2_ conditions. However, *in vivo* O_2_ gradients are highly dynamic, both spatially and temporally. The stepwise transitions and micro-gradients that EVTs experience cannot be fully reproduced *in vitro*, which may influence the observed lineage progression and functional phenotypes. Although we used 8% O_2_ in our WT characterization, most of our experiments, including scRNA-seq, were conducted at 2% and 21% O_2_, reflecting physiological levels during early placental development and the atmospheric levels commonly used in laboratories. A more detailed interrogation of trophoblast behaviour at intermediate O_2_ conditions, such as 8%, will be important to understanding the lineage progression toward the end of the first trimester.

Another limitation lies in the use of a global *HIF1α* knockout. While informative, this approach does not allow temporal control over HIF1α loss, preventing precise dissection of its stage-specific roles during trophoblast differentiation and maturation. Furthermore, we did not investigate the contribution of other HIF family members or alternative hypoxia-responsive pathways, which may also participate in O_2_ sensing and could account for the HIF1α-independent effects observed. Lastly, HIF1α protein and hypoxia pathway signatures detected at 21% atmospheric O_2_ may reflect microhypoxic niches created by diffusion limits in 3D cultures; intra-organoid O_2_ measurements could be required to fully separate O_2_ availability from HIF1α activity.

## ACKNOWLEDGMENTS

The work was supported by the Max Planck Gesellschaft (M.H., C.G.). We thank the Gerri lab (especially Hugo Raveton for project discussion, Subham Seal and Krista Briedis Gert for reviewing the manuscript), David Flores Benitez, Jacqueline Tabler for discussions and feedback on the manuscript. We thank the Tabler Lab for sharing reagents. We thank the Organoid and Stem Cell Facility (especially Christina Eugster-Oegema and Jula Peters for cell culture support), Light Microscopy Facility (especially Jan Peychl and Riccardo Maraspini for microscopy instructions), Scientific Computing (especially Lena Hersemann and Andre Gohr for discussions regarding scRNAseq experiments and raw data processing, Anthony Vega for developing image analysis pipelines and analyzing data and Ivey Sebastian for developing image analysis pipelines), Genome Engineering Facility (especially Mihail Sarov, Ilka Reichardt-Gomez, Julia Sigl for mutant design and generation), and the Protein Expression Purification and Characterization facility (Barbara Borgonovo for trouble shooting and discussions) at the Max Planck Institute of Cell Biology and Genetics for their outstanding support. This work was supported by the Dresden-concept Genome Center (DcGC) and scRNA-seq was partially carried out by Susanne Reinhardt and Josefine Kleinert. The DcGC and their employees are supported by the following sources: SMWK, BMFTR, DFG, CMCB, MPI-CBG, PLID.

## AUTHOR CONTRIBUTIONS

Conceptualization, J.L., C.G; data curation, J.L, C.G.; formal analysis, J.L., J.B., C.G.; investigation, J.L., J.B., M.B., O.E.; methodology, J.L., J.B., M.B., O.E.; visualization, J.L., C.G.; project administration, C.G.; funding acquisition and resources, M.H., C.G.; supervision, C.G.; writing – original draft, J.L., M.M., C.G; writing – review and editing, J.L., J.B., M.B., O.E., M.M., M.H., C.G.

## Supplementary figures

**Figure S1.**
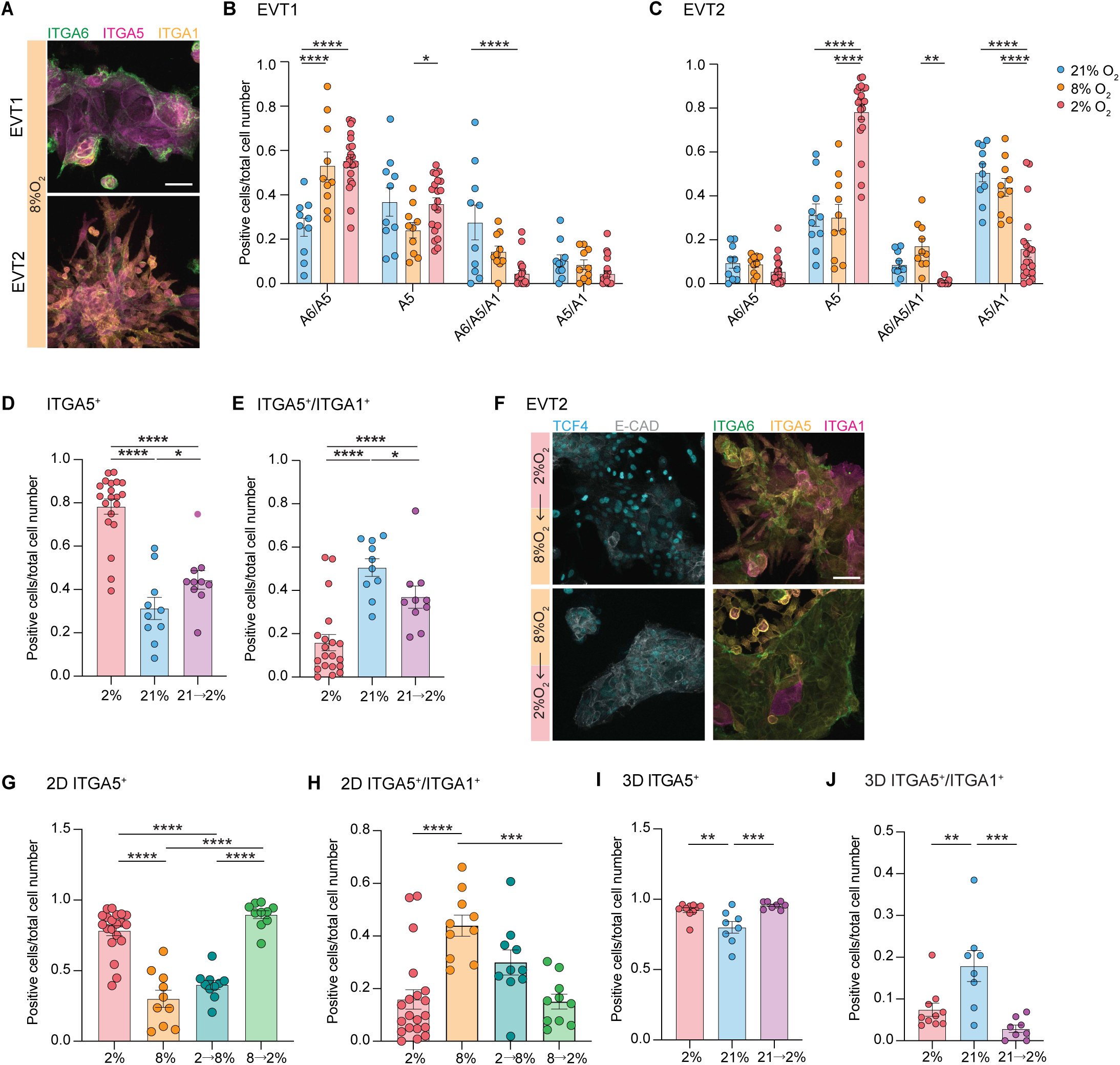
Integrin analysis reveals higher cell plasticity in trophoblasts at 8% O_2_ than 21% O_2_. (A) Immunofluorescence analysis of ITGA6, ITGA5, ITGA1 in EVT1 and EVT2 in WT EVTs at 8% O_2_. (B, C) Quantification of cell populations positive for ITGA6, ITGA5, and ITGA1 signal and combination of expression in EVT1 and EVT2 at 21%, 8% and 2% O_2_ (part of data also shown in Figure 1). (*n* = 10 cell colonies for 21% and 8%; *n* = 20 for 2%). (D) Quantification of ITGA5^+^ only, and ITGA5^+^/ITGA1^+^ populations in EVT2 in O_2_ switch experiment shown in Figure 1. (*n* = 10 cell colonies for 21% and switch experiment; *n* = 20 for 2%). (F) Immunofluorescence analysis of TCF4, E-CADHERIN, and ITGA6, ITGA5, ITGA1 in EVT2 in WT EVTs in O_2_ switch experiments. (G, H) Quantification of ITGA5^+^ only, and ITGA5^+^/ITGA1^+^ populations in EVT2 after O_2_ experiments shown in F. (part of data also shown in Figure 1) (*n* = 10 cell colonies for 8% and switch experiment; *n* = 20 for 2%). (I, J) Quantification of ITGA5^+^ only, and ITGA5^+^/ITGA1^+^ populations in EVT2 in switching experiment shown in Figure 2. (*n* = 10 organoids for 2% ; *n* = 8 for 21% and switch experiment). A5 = ITGA5; A6 = ITGA6; A1 = ITGA1. Error bars indicate mean +/- SEM. Statistical analysis by Two-way ANOVA and Tukey’s post-hoc test for multiple comparisons. * p<0.05; ** p<0.01; *** p<0.005; **** p<0.0005. Scale bars, 100 μm.

**Figure S2.**
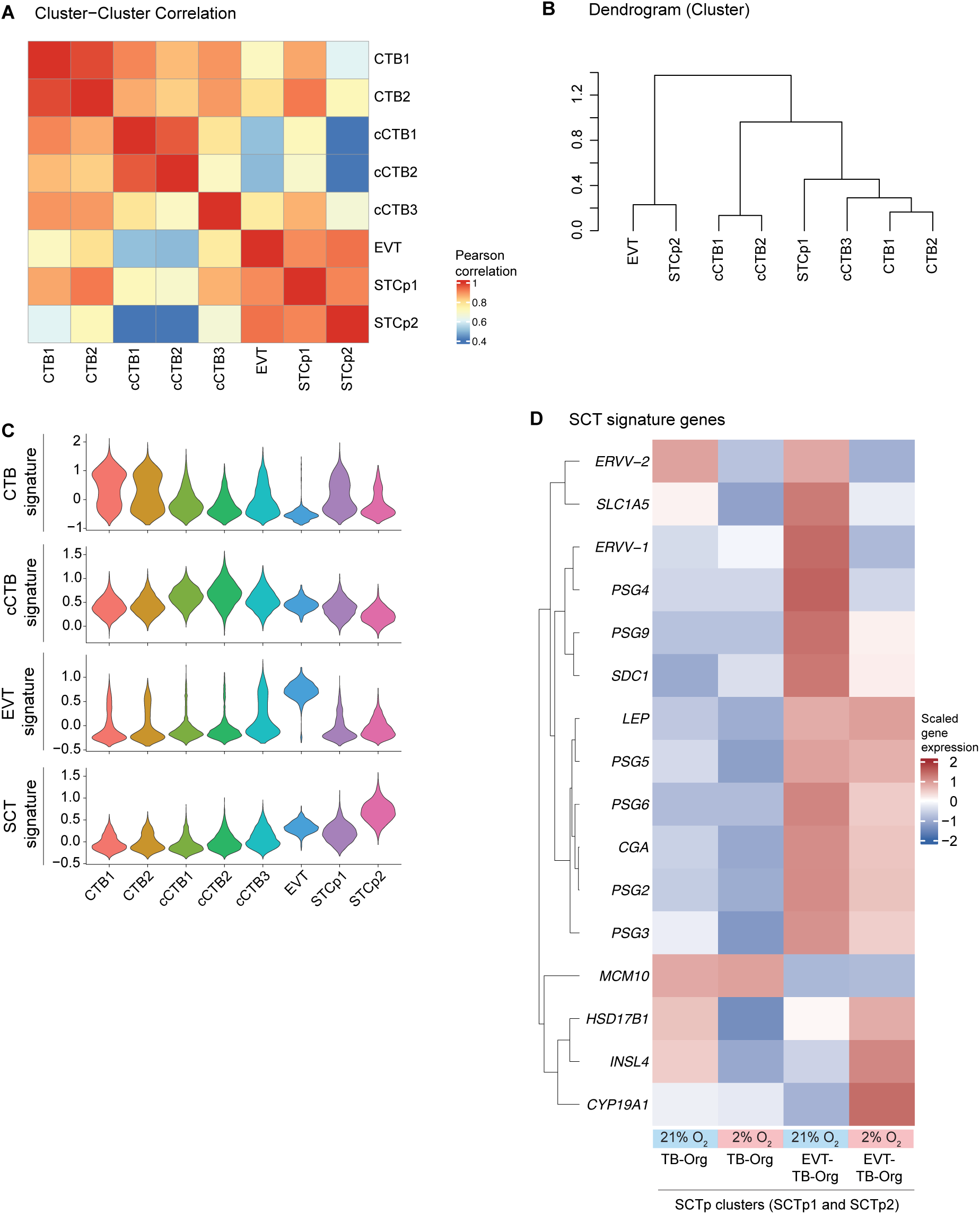
Transcriptional heterogeneity in trophoblast organoid model. (A) Pearson correlation across identified clusters in WT datasets. (B) Hierarchical clustering dendogram of clusters in WT datasets. (C) Violin plots of CTB, cCTB, EVT and SCT signature markers across clusters in WT datasets. Gene lists used for signature scores is provided in Supplementary Table S3. (D) Heatmap showing scaled gene expression of SCT signature genes in the SCTp clusters (SCTp) in WT TB-Orgs and EVT-TB-Orgs at 21% and 2% O_2_.

**Figure S3.**
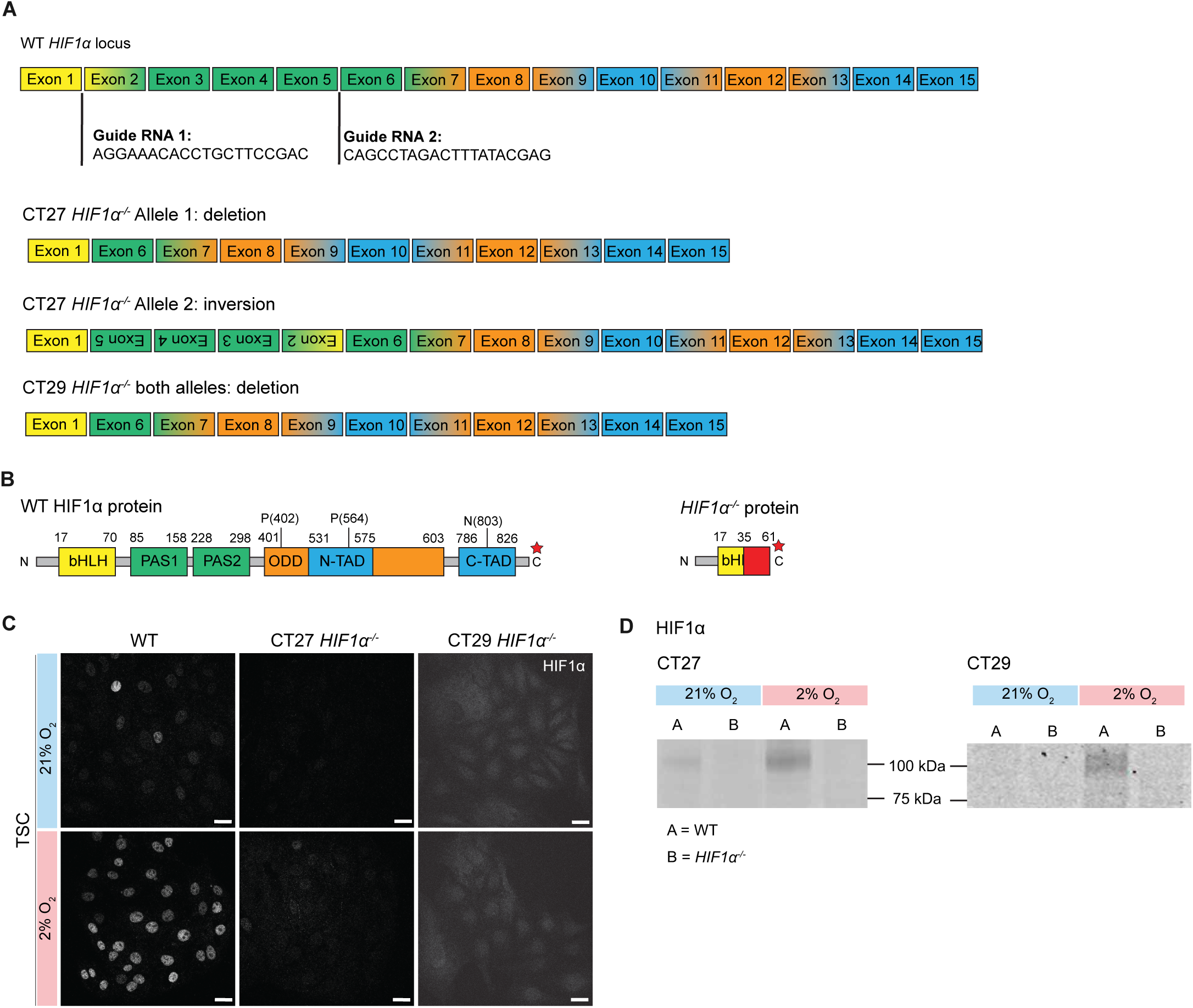
Generation strategy of *HIF1α^-/-^* trophoblast stem cell lines. (A) WT *HIF1α* locus with locations of guide RNAs, and *HIF1α^-/-^* CT27 and CT29 alelles. (B) WT HIF1α protein and predicted HIF1α^-/-^ mutant protein. Red stars show stop codon. bHLH, basic helix-loop-helix domain. PAS, PER-ARNT-SIM domains. ODD, O_2_-dependent degradation domain. N-TAD, N-terminal transactivation domain. C-TAD, C-terminal transactivation domain. (C) Immunofluorescence of HIF1α in WT and *HIF1α^-/-^* CT27 and CT29 TSCs at 21% and 2% O_2_. (D) Western blot of HIF1α in WT and *HIF1α^-/-^* CT27 and CT29 TSCs at 21% and 2% O_2_.

**Figure S4.**
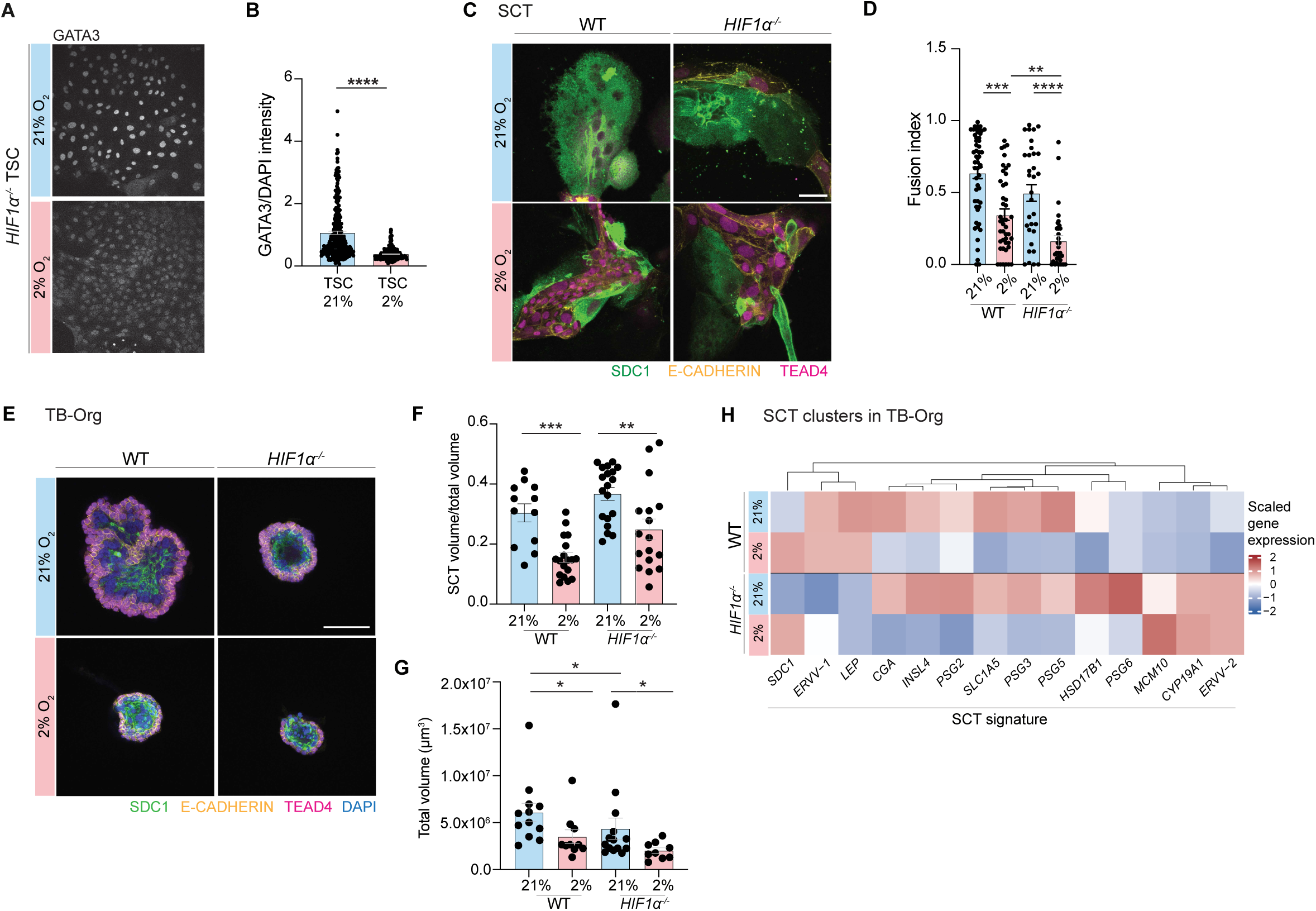
*HIF1α^-/-^* show reduced SCT formation at 2% O_2_. (A) Immunofluorescence of GATA3 in *HIF1α^-/-^* TSCs at 21% and 2% O_2_. (B) Quantification of nuclear GATA3 normalized fluorescence intensity in *HIF1α^-/-^* TSCs at 21% and 2% O_2_. (*n* = 266 cells for 21%; *n* = 241 for 2%). Statistical analysis by Mann-Whitney U test. (C) Immunofluorescence images of E-CADHERIN, TEAD4, SDC1 in WT and *HIF1α^-/-^* SCTs at 21% and 2% O_2_. (D) Quantification of fusion index in WT and *HIF1α^-/-^* SCTs at 21% and 2% O_2_ (WT data also shown in Figure 1). (*n* = 53 cell colonies for WT 21%; *n* = 44 for WT 2%; *n* = 32 for *HIF1α^-/-^* 21%; *n* = 32 for *HIF1α^-/-^*2%.) Statistical analysis by Two-way ANOVA and Tukey’s post-hoc test for multiple comparison. (E) Immunofluorescence images of E-CADHERIN, TEAD4, SDC1, and DAPI signal in WT and *HIF1α^-/-^* TB-Orgs at 21% and 2% O_2_. (F) Quantification of SCT volume over total organoid volume in WT and *HIF1α^-/-^* TB-Orgs at 21% and 2% O_2_ (WT data also shown in Figure 2). (*n* = 12 organoids for WT 21%; *n* = 18 for WT 2%; *n* = 19 for *HIF1α^-/-^* 21%; *n* = 17 for *HIF1α^-/-^* 2%). Statistical analysis by Two-way ANOVA and Tukey’s post-hoc test for multiple comparison. (G) Quantification of total organoid volume in WT and *HIF1α^-/-^* TB-Orgs at 21% and 2% O_2_ (WT data also shown in Figure 2). (*n* = 12 organoids for WT 21%; *n* = 10 for WT 2%; *n* = 14 for *HIF1α^-/-^* 21%; *n* = 9 for *HIF1α^-/-^* 2%). Statistical analysis by Two-way ANOVA and Tukey’s post-hoc test for multiple comparison. (H) Heatmap of scaled gene expression of SCT signature in SCTp clusters in WT and *HIF1α^-/-^* TB-Orgs at 21% and 2% O_2_.

**Figure S5.**
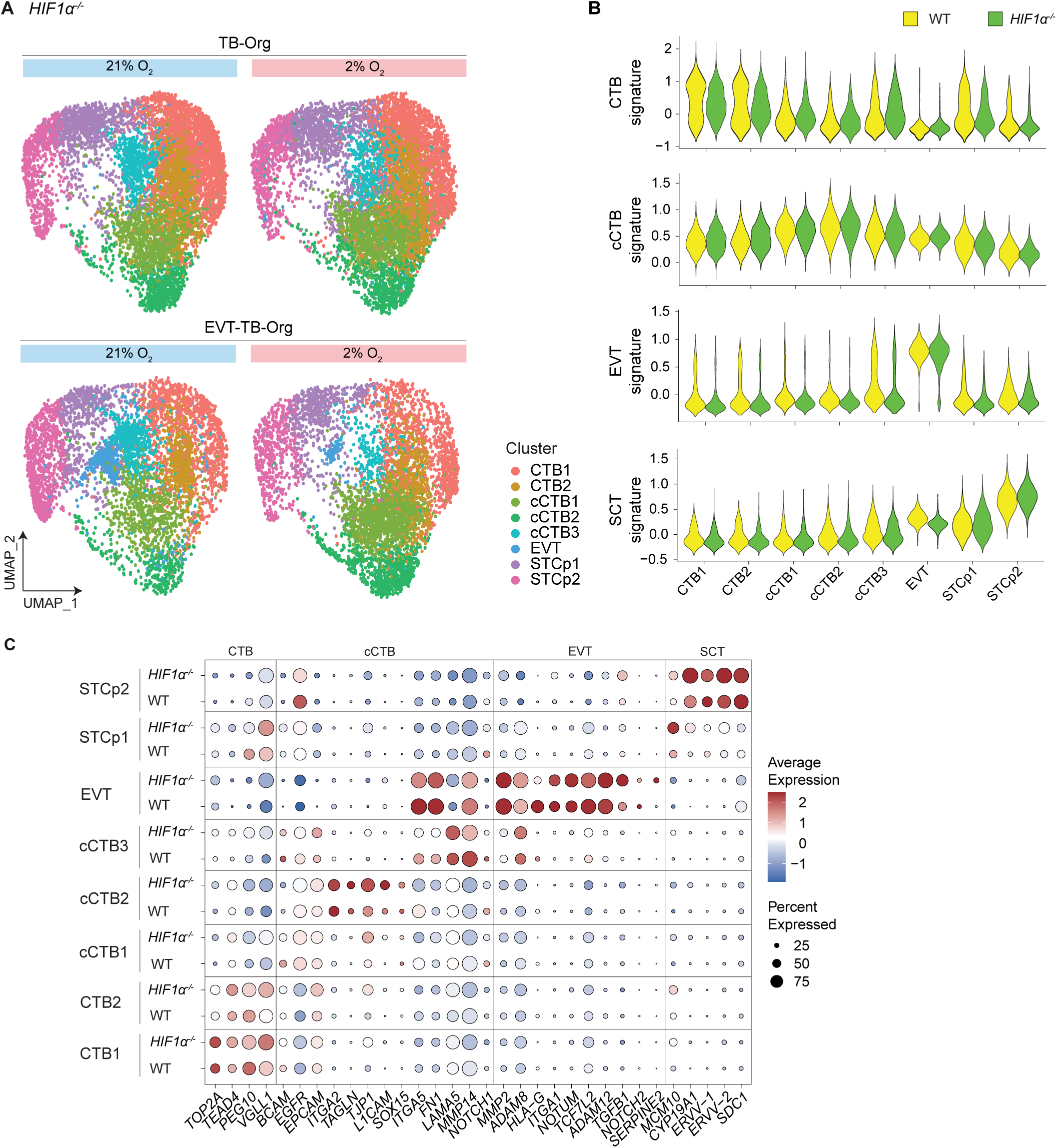
Comparative analysis of WT and HIF1α^-/-^ trophoblast organoids across O_2_ conditions. (A) UMAP of *HIF1α^-/-^* trophoblast organoids split by condition, showing TB-Orgs and EVT-TB-Orgs at 21% and 2% O₂ and colored by cell type. Compare with Figure 3B for the UMAP of WT organoids. (B) Violin plots of CTB, cCTB, EVT and SCT signature markers across identified clusters in WT and *HIF1α^-/-^* datasets. (C) Dot-plot showing gene expression of representative signature markers for cluster and genotype (WT and HIF1α^-/-^). The dot size corresponds to the ratio of cells expressing the gene in the population and the colour scale corresponds to the cell type averaged gene expression level.

## Supplementary Table legends

**Supplementary Table S1. List of reagents and antibodies used.**

List of reagents for cell culture, Western Blot and single cell RNA sequencing used, and list of primary and secondary antibodies used for Western blot and immunofluorescence.

**Supplementary Table S2. Off-target screening in *HIF1α^-/-^* hTSC lines**.

Sequencing results of the 5 top off-target sites for each guide RNA.

**Supplementary Table S3. Single-cell analysis of trophoblast-derived organoids.**

Signature genelists for CTB, cCTB, SCT and EVT population annotation; Differentially expressed genes per cluster in WT trophoblast organoids (log2fc>0.25, min.pct=0.1, wilcoxon-test); Differentially expressed genes per cluster and genotype in trophoblast organoids (log2fc>0.25, min.pct=0.1, wilcoxon-test)

**Supplementary Table S4. scRNAseq comparison between WT and *HIF1α^-/-^* normoxia.**

Differentially expressed genes in cCTB1, cCTB2, cCTB3 and EVT (WT vs *HIF1α^-/-^*) in EVT-TB organoids in normoxia conditions (21%) (log2fc>0.25, min.pct=0.1, p-adj-value < 0.05, wilcoxon-test)

**Supplementary Table S5. Gene enrichment analysis per population (WT vs *HIF1α^-/-^*).**

Hallmark database (MSigDB_Hallmark 2020) gene enrichment analysis of upregulated genes in cCTB1, cCTB2, cCTB3 and EVT (WT vs *HIF1α^-/-^*) in EVT-TB organoids in normoxia condition (21% O2).

## METHODS

### Cell lines

CT27 (RCB4936; female) and CT29 (RCB4937; male) patient-derived trophoblast stem cell (TSC) lines used in this study were obtained from the RIKEN stem cell bank (https://cellbank.brc.riken.jp/cell_bank/CellInfo/?cellNo=RCB4936; https://cellbank.brc.riken.jp/cell_bank/CellInfo/?cellNo=RCB4937&lang=En) after approval from the Robert Koch Institute. The CT27 TSCs were derived from placental CTB cells as described in ref. ^62^. Human placentas were obtained from healthy women with signed informed consent of the donors, and the approval of the Ethics Committee of Tohoku University School of Medicine (Research license 2014-1-879).

### Human trophoblast stem cell culture and differentiation

The human trophoblast stem cell lines CT27 and CT29 were cultured similar as previously described^41^. All used cell culture reagents can be found in **Supplementary Table S1**. A 6-well plate was pre-coated with human 5µg/ml Col IV for 10min at 37°C. Cells were seeded at a density of 4 x 10^4^per well and cultured in 2ml of hTSC medium containing 10µM Y2763 over night and media was changed the next day to regular hTSC medium (DMEM/F12 supplemented with 0.1 mM 2-mercaptoethanol, 0.2% FBS, 0.5% Penicillin-Streptomycin, 0.3% BSA, 1% ITS-X supplement, 1.5 mg/ml L- ascorbic acid, 50 ng/ml EGF, 2 mM CHIR99021, 0.5 mM A83-01, 1 mM SB431542, 0.8 mM VPA and 5 µM Y27632). Cells were maintained at 37°C, 21% O_2_ and 5% CO_2_ and media was changed every other day until cells reached 80-90% confluency. At 80-90% confluency (usually after 3-4 days) cells were rinsed with PBS and then dissociated with TrypLE for 10-15min at 37°C, finally passaged to a new Col IV-coated 6-well plate. Cells for experiments were used between passages 5-35.

For differentiation into STB as previously described^41^ cells were seeded at a density of 1 x 10^5^on a 6-well plate pre-coated with 2,5µg/ml Col IV for Western blot analysis, and for immunofluorescence analysis at a density of 1,5 x 10^4^ on an 8-well IBIDI dish pre-coated with 2,5µg/ml Col IV. Cells were cultured in 2ml or 400µl STB (DMEM/F12 supplemented with 0.1 mM 2-mercaptoethanol, 0.5% Penicillin-Streptomycin, 0.3% BSA, 1% ITS-X supplement, 2.5 mM Y27632, 50 ng/ml EGF, 2 mM forskolin, and 4% KSR) medium for 72h to 96h replacing media after 2 days.

For the induction of EVT differentiation as previously described^41^ on day 0 cells were suspended in EVT1 medium (DMEM/F12 supplemented with 0.1 mM 2-mercaptoethanol, 0.5% Penicillin-Streptomycin, 0.3% BSA, 1% ITS-X supplement, 100 ng/ml NRG1, 7.5 mM A83-01, 2.5 mM Y27632, and 4% KSR, and 2% Matrigel) and seeded at a density of 9 x 10^4^ on a 6-well plate pre-coated with 1µg/ml Col IV. To asses timepoints day 2 (EVT1) and day 4 (EVT2) of differentiation cells were seeded at a density of 0,75 x 10^4^ in an 8-well IBIDI dish pre-coated with 1µg/ml Col IV and fixed at the respective time points. Cells were cultured in EVT1 medium for two days. On day 2 media was exchanged for EVT2 medium (DMEM/F12 supplemented with 0.1 mM 2-mercaptoethanol, 0.5% Penicillin-Streptomycin, 0.3% BSA, 1% ITS-X supplement, 7.5 mM A83-01, 2.5 mM Y27632, and 4% KSR, and 0,5% Matrigel) for another two days. On day 4 cells from 6-well plate were dissociated with TrypLE for 10-15min at 37°C, passaged to a pre-coated 8-well IBIDI at a split ratio of 1:20, and cultured for another 2 days in EVT3 medium (DMEM/F12 supplemented with 0.1 mM 2-mercaptoethanol, 0.5% Penicillin-Streptomycin, 0.3% BSA, 1% ITS-X supplement, 7.5 mM A83-01, 2.5 mM Y27632, and 0,5% Matrigel).

For assessment of cells at 2% or 8% O_2_ concentration all media and solutions were equilibrated at 2% or 8% O_2_ respectively prior to administration, and cells were handled in the hypoxia station (Whitley H35 HEPA hypoxystation).

### TB-Org generation and EVT-TB-Org differentiation

TB-Orgs were generated as previously described^45,46^ from hTSCs grown in hTSC medium. When cells reached 80% confluency they were dissociated using TrypLE for 10-15min at 37°C, resuspended in TOM (Advanced DMEM/F12, N2 supplement final conc. 1x, B27 supplement minus vitamin A at a final conc. of 1x, Primocin 100 μg/mL, N-Acetyl-L-cysteine 1.25 mM, L-glutamine 2 mM, recombinant human EGF 50 ng/mL, CHIR99021 1.5 μM, recombinant human R-spondin-1 80 ng/mL, recombinant human FGF-2 100 ng/mL, recombinant human HGF 50 ng/mL, A83-01 500 nM, prostaglandin E2 2.5 μM, Y-27632 2 μM) at a final cell number of 5.000 cells per 100µl Matrigel. 12,5µl drops were placed in each well of an 8-well IBIDI dish for immunofluorescence and 25µl drops in a 12 well plate for Western Blot analysis or scRNAseq. After one minute of benchtop incubation, dishes were turned upside down, incubated at 37°C for at least 15min to solidify, then 400µl of TOM were added to each well and cultured for 7 days until fixation for analysis of syncytium formation. For differentiation into EVT-TB-Orgs differentiation we followed the previously published protocol from Sheridan et al.^66^ EVT-TB-Org differentiation was initiated on day 5 after TB-Org set-up by changing media to EVT1 medium (Advanced DMEM/F12, L-Glutamax 2 mM, 2-mercaptoethanol 0.1 mM, penicillin/streptomycin solution 0.5%, BSA 0.3%, ITS-X supplement 1%, NRG1 100 ng/mL, A83-01 7.5 μM, KSR 4%) and culturing EVT-TB-Orgs for 2-4 days. Then media was exchanged to EVT2 medium (Advanced DMEM/F12, L-Glutamax 2 mM, 2-mercaptoethanol 0.1 mM, penicillin/streptomycin solution 0.5%, BSA 0.3%, ITS-X supplement 1%, A83-01 7.5 μM, KSR 4%) for another 2-3 days. For time sensitive experience as time course in IF images and rescue experiments by O_2_ shifts, we always used the exact same differentiation time of 2 days for EVT1 medium and 2 days for EVT2 medium. For all others we followed the Sheridan et al. protocol of tracking the degree of differentiation, always treating genotype conditions equally at the same time for comparability.

For assessment of cells at 2% or 8% O_2_ concentration all media and solutions were equilibrated at 2% or 8% O_2_ respectively prior to administration, and cells were handled in the hypoxia station (Whitley H35 HEPA hypoxystation).

### Generation of *HIF1α* knockout in hTSC CT27 and CT29

*HIF1α* knock-out was generated through deletion of critical exons 2-5 (ca. 7 kb). Deletion of these exons generates an out-of-frame transcript with a premature stop codon which leads to a truncated protein of 39 aa.

Guide RNAs specific to human *HIF1α* locus were selected based on low off-target activity using http://crispor.tefor.net. The guide RNAs were ordered as crRNA from Integrated DNA Technologies (IDT), upstream guide RNA 5’-AGGAAACACCTGCTTCCGAC and downstream guide RNA 5’-CAGCCTAGACTTTATACGAG.

2 x10^5^ hTSC CT27 were transfected with 31 pmol, and CT29 with 18pmol Cas9 protein (IDT Cat.no. 1081061) complexed with 46 pmol (CT27) or 22pmol (CT29) crRNA (IDT, Alt-R®) and tracrRNA (IDT Cat.no. 1072534) using the Neon electroporator device (1300 V, 20 ms, 1 pulse) and kits (ThermoFisher MPK1096) according to the manufacturer instructions. Electroporated cells were plated as described above into hTSC medium with 10 µM Y2763. 24h post-transfection medium was changed to hTSC medium containing 5µM Y2763. 72 h post-transfection cells were single-cell sorted into 96-well plates containing 100 µl hTSC medium with 10 µM Y2763. Cell sorting was performed in a BD FACSAria Fusion flow cytometer (Beckton Dickinson). 48h post-sorting medium was changed to hTSC medium containing 5µM Y2763. Single-cell clones were expanded from 96-well about 10-12 days post-sorting. Genotyping samples were taken during splitting and genotyping was conducted by PCR. Genomic DNA was extracted using the QuickExtract DNA extraction kit (Epicentre) following the manufacturer’s instructions. PCR was performed using Phusion Flash High-Fidelity PCR Master Mix (ThermoFisher) with gene-specific primers spanning the deletion, forward primer 5’-GGAAGACAGCTTTTGCTTGGT and reverse primer 5’-AATCAGCACCAAGCAGGTCA (product size for *wild-type* is 7970 bp and 880 bp for the knockout). Amplicons of the deletion alleles were verified by Sanger Sequencing. To exclude remaining wild-type allele, flanking PCR with one primer outside and one primer inside the deletion was performed. For the 5’ *wild-type* flanking PCR we used 5’-GGAAGACAGCTTTTGCTTGGT forward primer and 5’-TCAAAACATTGCGACCACCTTC reverse primer (product size for *wild-type* is 1085 bp and no product for the knockout), and for 3’ *wildtype* flanking PCR we used 5’-5’-GTAATTTTCTGCCTGCTTTACTAGA forward primer and 5’- AATCAGCACCAAGCAGGTCA reverse primer (product size for *wild-type* is 1365 bp and no product for the knockout). The five most common off targets for each crRNA were screened using PCR and Sanger sequencing the exact primer pairs are listed in **Supplementary Table S2**.

### Immunofluorescence staining, imaging and image analysis

For imaging 2D hTSCs, TB-Orgs and EVT-TB-Orgs in IBIDI dishes were treated equally. Samples were rinsed with PBS prior to fixation in 4% PFA for 15min at RT. Samples cultured at 2% O_2_ were handled in the hypoxia station with equilibrated PBS until PFA was added. After fixation samples were rinsed 3x with PBS. Samples were permeabilized at RT for 10 min with wash buffer (PBS containing 0,5% Tween 20), then blocked at RT for 30min in blocking solution (PBS containing 0,5% Tween 20 and 3% BSA). Primary antibodies were diluted in blocking solution and incubated over night at 4°C. Samples were washed in wash buffer by rinsing 3x and washing at least 2x for 5min at RT. Secondary antibodies were diluted in blocking solution and incubated for 2h at RT together with DAPI and Phalloidin-660. Samples were washed in washed in wash buffer by rinsing 3x and washing at least 2x for 5min at RT. For clearing, samples were rinsed 2x with PBS and then kept in Glycerol-Fructose solution^67^. Confocal imaging was done using a multiphoton ZEISS LSM 780 NLO inverted system with a Zeiss LD LCI Plan-Apo 25x (0.8 NA) multi-immersion DIC objective. Brightfield images were taken on a Nikon Eclipse Ts2-LS with a 10x phase contrast objective.

2D syncytium analysis was done by determining the fusion index. STB differentiated cells were stained for syndecan-1, E-cadherin, TEAD4 and DAPI. The confocal z-stack images were max projected in FIJI (), and areas positive for syndecan-1 and negative for E-cadherin and TEAD4 were manually encircled and nuclei were counted by manually selecting them. Fusion index was calculated as previously described^23^ by subtracting the number of syncytia detected from the number of nuclei in syncytium and then dividing by the total number of nuclei. TEAD4 positive cells were quantified in the same images by subtracting nuclei in syncytium from the total number of nuclei.

Nuclear intensity quantification in 2D. TSCs, and EVTs were stained for GATA3, TCF4 and DAPI. For image quantification of TCF4 and GATA3 in 2D, images were processed using a custom Arivis Pro 4.3.1 pipeline. Images were denoised to reduce background noise, and nuclei were segmented based on DAPI staining using a blob detection approach. The resulting nuclear masks were used to quantify the intensity of TCF4 and GATA3 within each nucleus relative to DAPI. To exclude debris and over-segmented objects, nuclei were filtered based on area. Objects smaller than 700 µm^3^ were removed from all analyses. This filtering step was applied individually to each replicate image.

For 2D EVT integrin classification, EVT differentiated cells were stained for ITGA1, ITGA5, ITGA6 and DAPI. Image analysis was performed on CentOS Linux 7.4.1708 running Python 3.10.7. We used a pretrained 2D Cellpose model (‘nuclei’), and applied it to maximum projections of our data. We labeled a subset of nuclei segmentation data using manually selected intensity thresholds, which was the same for each integrin subtype and used a multi-class logistic regression model (scikit-learn, Logistic Regression) with 5-fold cross-validation for classification (0.978 +- 0.014).

For the 3D Volume quantification TB-Orgs were stained for syndecan-1, E-cadherin, TEAD4 and DAPI. TB-Org syncytium image analysis was performed on CentOS Linux 7.4.1708 running Python 3.10.7. For TB-Org segmentation, for each image volume we first applied a Gaussian filter (sigma=5) and then used the Otsu threshold from the scikit-image library to obtain an initial organoid mask. We then kept only the largest thresholded volume and applied post-processing steps of binary closing and erosion to further refine the mask. We used the triangle threshold to segment syncytium signal within the organoid and calculated the fraction of the organoid volume occupied by this signal.

Nuclear intensity quantification in EVT-TB-Orgs. EVT-TB-Orgs were stained for at least TCF4 and DAPI. Fluorescence images were acquired as CZI files using confocal microscopy. Images consisted of DAPI (nuclei) and TCF4 immunofluorescence channels. Raw data were imported in Python using aicspylibczi. For each image, the middle z-plane was extracted for analysis. Nuclear segmentation was performed on the DAPI channel using Cellpose 2.0^68^ with the built-in nuclei model and a fixed diameter of 20 pixels. Segmentation masks were visually inspected but not manually corrected. Nuclear area was obtained directly from Cellpose output. TCF4 intensity quantification was performed by applying the DAPI-derived segmentation masks to the TCF4 channel. For each nucleus, we calculated mean nuclear TCF4 intensity, DAPI intensity, and the TCF4/DAPI ratio. Background was estimated as the median non-nuclear intensity per image and subtracted before quantification. To ensure data quality, nuclei smaller than 300 µm² were excluded. All measurements were exported with their corresponding source image identifiers. Replicates belonging to the same condition were aggregated.

For EVT-TB-Org integrin classification, EVT-TB-Orgs were stained for ITGA1, ITGA5, ITGA6 and DAPI. Image analysis was performed on CentOS Linux 7.4.1708 running Python 3.10.7. We used a pretrained 2D Cellpose model (‘nuclei’), and applied it to the middle slice of the z-stack image. We labeled a subset of nuclei segmentation data using manually selected intensity thresholds, which was the same for each integrin subtype and used a multi-class logistic regression model (scikit-learn, Logistic Regression) with 5-fold cross-validation for classification (0.978 +- 0.014).

### Protein extraction and western Blotting

Proteins were extracted from EVT-TB-Org and TB-Org by releasing them first from Matrigel by suspending drops of Matrigel in ice cold recovery solution for 15min, and from 2D culture after stem cells reached 80-90% confluency or at the indicated time points of differentiation by washing cells with ice cold PBS and then scraping cells from the plate in ice cold RIPA Buffer, samples were centrifuged, supernatant was collected for BCA assay and set aside and the sample was directly boiled for 5min in 1x Laemmli buffer at 95°C. BCA was done using a kit from Thermo Fisher according to the manufacturer instructions. Samples were treated with benzonase (according to manufacturers instructions), then 10-30µg of protein were loaded on the SDS gel, and transferred onto nitrocellulose membranes. Membranes were blocked for 2h at RT in blocking solution (4% BSA and 0,4% Tween 20 in tris buffered saline TBST), incubated in primary antibody solution ON at 4°C, washed with TBST and incubated in secondary antibody solution for 1h at RT. For visualization LI-COR Odyssey Sa Infrared-Imaging-System was used. Quantification was done using LI-COR ImageStudio software. For reagents and antibodies used see details in **Supplementary Table S1**.

### TB-Org collection a fixed RNA profiling with 10x Genomics

Single Cell suspension was prepared following 10x Genomics protocol for Chromium Fixed RNA Profiling by releasing EVT-TB-Orgs or TB-Orgs from Matrigel by incubating in ice cold cell recovery solution for 40min, washing with ice cold PBS, incubating in TrypLE with DNAseI at 37°C for 7min vortexing samples in the middle of incubation time. If dissociation wasn’t satisfactory incubation with TrypLE was prolonged for max 2min more. To finally singularize cells, appropriate media (EVT2 medium or TOM) was added to stop reaction and pipetting up and down to generate single cell suspension. Cells were strained using 40µm filter and counted. Fixation and storage was done using the 10x Genomics kit (PN-1000414) according to the manufacturer instructions. In short, samples were incubated in formaldehyde-containing fixation buffer at 4 °C for 22 hours. The fixation was terminated by replacing the fixation buffer with a quenching buffer. After determining the cell concentration, fixed samples were prepared for long-term storage. Therefore, a pre-warmed enhancer was added at a 1:10 ratio to fixed cells resuspended in a quenching buffer. Finally, 50 % glycerol was added to this mixture for a final concentration of 10 % and stored at – 80°C for up to six months after fixation.

Thawing and subsequent processing of the fixed samples followed the 10x Genomics user manual guidelines for Chromium fixed RNA profiling and using the Chromium Fixed RNA Kit, Human Transcriptome (PN-1000476). In brief, the thawed samples were washed in 0.5X PBS + 0.02 % BSA and counted manually using a hemocytometer. 12 samples with more than 0.5 million cells were multiplexed by hybridization with sample-specific barcode-containing gene probes at 42 °C overnight for 21 hours to profile approximately 18000 human genes. After stopping the hybridization and washing, samples were individually counted before pooling to achieve an even representation of samples. Cell concentration was adjusted after extensive washing and filtering through a 30 µm cell strainer. Sample pools were mixed with reverse transcription reagents before loading 150.3k pooled cells or nuclei per lane of a Chromium Single Cell Q Chip^69^ from Chromium Next GEM Chip Q Single Cell Kit (PN-1000422) and therefore targeting an output of 96k cells. After gel bead in emulsion (GEM) generation, sample-barcoded probes were ligated to barcode-containing oligo sequences and immediately subjected to reverse transcription. Generated cDNA was recovered from GEMs and pre-amplified for eight cycles. Prior to library construction, amplified cDNA underwent 1.8× purification using SPRIselect beads (Beckman Coulter, B23319). Based on the total number of loaded cells and the cell types, 9 cycles were used for fixed RNA library construction with a sample index PCR to add TS indices. After 1.0× purification using SPRIselect beads, final libraries underwent quality control using the Fragment Analyzer (Agilent Technologies, Inc.), aiming to verify a 270 base pair peak. Fixed RNA libraries were sequenced on S4 flow cell using the Illumina NovaSeq 6000 sequencing system in paired-end mode (R1/R2: 100 cycles, I1/I2: 10 cycles), aiming to reach ∼10000 reads per cell on average.

### Extension of gene panel 10X Chromium Single Cell Gene Expression Flex

The default human gene panel provided by 10x Genomics for the Chromium Single Cell Gene Expression Flex (Fixed RNA Profiling) assay was used as the basis for transcriptome profiling. Because several classical HLA class I genes are absent from this panel, it was extended with custom-designed probes targeting *HLA-A*, *HLA-B*, *HLA-C*, *HLA-E*, and *HLA-G*. Probe design followed the recommendations outlined in the 10x Genomics Technical Note (CG000621) and was further refined using a custom computational pipeline developed for this study. To identify candidate probe regions, publicly available scRNA-seq data (GEO: GSE174481) generated from human trophoblast stem cell–derived organoids (cell lines C27 and C29) before and after EVT differentiation was used. Reads were mapped as bulk RNA-seq using Hisat2 v2.2.1 within the nf-core/rnaseq v3.14.0 workflow to the Homo sapiens GRCh38 reference genome (Ensembl release 111). Candidate HLA gene segments were identified through visual inspection in IGV, selecting regions ≥60 nt that showed consistent read coverage and were free of SNPs across all four samples.

Next, a custom off-target detection workflow was implemented to assess probe specificity. A Bowtie v1.3.1 index was generated from all coding and non-coding sequences of the GRCh38 genome. For each candidate region, the left-hand side (LHS) and right-hand side (RHS) sequences of putative probes were mapped independently to the index, allowing up to three mismatches. Probes were retained only if the following criteria were met: (i) LHS and RHS did not map adjacently to any off-target loci; (ii) both fragments exhibited GC content between 44–72%; (iii) the LHS terminated with a thymine at the 3′ end; and (iv) LHS and RHS did not overlap when mapped to the intended target gene. Using these criteria, two probes each were selected for HLA-B, HLA-C, HLA-E, and HLA-G, and one probe for HLA-A, which were incorporated into the extended gene panel.

### scRNA-seq sample demultiplexing with Cellranger

Reads generated per batch using the 10x Genomics Chromium Single Cell Gene Expression Flex (Fixed RNA Profiling) assay were demultiplexed with Cell Ranger v9.0.1. The human reference dataset (refdata-gex-GRCh38-2024-A) provided by 10x Genomics was used. Prior to running Cell Ranger, this reference was modified by extending the Chromium Human Transcriptome Probe Set v1.1.0 (0_GRCh38-2024-A) with the custom HLA gene probes described above.

### scRNA-seq data preprocessing and quality control

Individual single-cell (SC) samples were further processed and filtered using custom scripts implemented in R v4.0.5, Seurat v4.1.1^70^, and SCTransform v0.3.4^71^. For each sample, the following filtering steps were applied: (i) non–protein-coding genes and genes detected in fewer than 250 cells were removed; (ii) cells with fewer than 750 total read counts or fewer than 500 detected genes were excluded; and (iii) cells were removed if more than 15% of reads mapped to mitochondrial genes. All HLA genes were retained provided that at least one read was detected. Data normalization was performed using SCTransform without correction for additional confounding variables.

SCT-transformed dataset integration was performed using the default parameters of Seurat’s SCT integration workflow (https://satijalab.org/seurat/archive/v4.3/sctransform_v2_vignette). In short, integration features (3,000 genes) were selected using *SelectIntegrationFeatures* function, integration anchors were identified using *FindIntegrationAnchors* (normalization.method = “SCT”, reduction =”cca”) and the datasets were integrated using *IntegrateData* using SCT normalization. Following integration, dimensionality reduction was performed by computing the first 30 principal components (PCs). The neighborhood graph was constructed using the Seurat built-in function *FindNeighbors* (dims = 1:30), and clustering was performed using the Louvain algorithm with a resolution of 0.1. Following integration, a second layer of quality control was applied. Clusters were identified as putative low-quality if they satisfied at least two of the following criteria: elevated mitochondrial read fraction, low library complexity (low total counts and detected genes), or high expression of immediate early genes^72,73^. These low-quality clusters were removed, and dimensionality reduction, neighborhood graph construction, clustering, and UMAP embedding were repeated. After this additional filtering step, the final dataset comprised 50,390 high-quality cells.

Additionally, data were normalized to counts-per-ten-thousand and then logp1-transformed using LogNormalize followed by ScaleData, which was employed for computing cluster markers (FindAllMarkers, min.pct = 0.1, only.pos = TRUE, logfc.threshold = 0.25), for visualization of gene expression and for any other downstream analysis. Marker genes for each cell type are provided in **Supplementary Table S3** (log2FC > 0.25, adjusted p < 0.05).

Cell state annotation was performed using canonical lineage-specific marker genes: cytotrophoblasts (CTB: *TOP2A*, *TEAD4*, *PEG10*, *VGLL1*), columnar CTB (cCTB: *ITGA2*, *TJP1*, *SOX15*, *L1CAM*, *ITGA5*), extravillous trophoblasts (EVT: *MMP2*, *HLA-G*, *ITGA1*, *NOTUM*, *ADAM12*), and syncytiotrophoblasts (SCT: *CYP19A1*, *ERVV-1*, *ERVV-2*, *INSL4*). Cell type-specific gene signatures^4,5,74^ were constructed and annotation was validated using the *AddModuleScore* built-in function in Seurat. Full gene signature lists are provided in **Supplementary Table S3**. Following annotation, average gene expression per cluster was calculated using the *AverageExpression* function (assay = "RNA", slot = "data"), and Pearson correlation coefficients were computed between cluster-average profiles to generate a cluster–cluster correlation matrix. Correlation matrices and dendrograms were visualized to support the assessment of annotated cluster relationships.

Differential gene expression analysis between wild-type and HIF1A knockout cells was performed for each population of EVT-TB organoids under both hypoxic (2% O₂) and normoxic (21% O₂) conditions using the FindMarkers function (FindAllMarkers, min.pct = 0.1, logfc.threshold = 0.25). Functional enrichment analysis was conducted using the EnrichR web tool via the enrichR R package (v3.2). Population-specific DEGs were tested for enrichment in MSigDB Hallmark gene sets, and terms with an adjusted p-value < 0.05 were considered significant. The complete list of population-specific DEGs and the results of the MSigDB Hallmark gene set analysis are provided in **Supplementary Table S4 and S5,** respectively.

Pathway activity was inferred using PROGENy (v.1.17.1)^49^ as implemented in the decoupleR package and as described above for the refined ductal organoids. In short, PROGENy scores were computed on the normalized gene expression counts with parameters set to “top=500 target genes, perm=1, and organism=”Human”). The resulting hypoxia pathway activity matrix was scaled and summarized by calculating the average activity per cell type and condition. The HIF1A regulon was retrieved using the get_collectri function from the decoupleR package (v2.5.3) with the following parameters organism = "human" and split_complexes = FALSE. Entries were filtered to retain only those where the source was HIF1A and the mode of regulation (mor) equaled 1. Glycolysis and oxidative phosphorylation gene sets were obtained from the corresponding HALLMARK gene lists using the msigdbr package (v25.1.1). Gene set activity scores were calculated for each cell using the *AddModuleScore* built-in function in Seurat.

Single-cell data visualization was done using built-in Seurat functions, scCustomize (v.1.1.3), pheatmap (v1.0.12) and ggplot (v.3.4.2).

### Statistical analysis

Data are reported as mean values with standard error of the mean (SEM). Analyses were carried out using GraphPad Prism (version 9.3.1). For single comparisons non-parametric Mann-Whitney Tests were performed. For multiple comparisons two-way ANOVA with Tukey correction was performed.

## Data availability

The scRNA-seq datasets generated during this study are available at the Gene Expression Omnibus (GEO) under accession numbers GSE312669.

## REFERENCES

1 Velicky, P. et al. Genome amplification and cellular senescence are hallmarks of human placenta development. PLoS genetics 14, e1007698 (2018). 10.1371/journal.pgen.1007698

2 Gauster, M., Moser, G., Wernitznig, S., Kupper, N. & Huppertz, B. Early human trophoblast development: from morphology to function. Cell Mol Life Sci 79, 345 (2022). 10.1007/s00018-022-04377-0

3 Meinhardt, G. & Haider, S. NOTCH, WNT, and TGFβ: Key pathways in extravillous trophoblast formation and differentiation. Placenta (2025). 10.1016/j.placenta.2025.07.082

4 Shannon, M. J. et al. Cell trajectory modeling identifies a primitive trophoblast state defined by BCAM enrichment. Development 149 (2022). 10.1242/dev.199840

5 Shannon, M. J. et al. Single-cell assessment of primary and stem cell-derived human trophoblast organoids as placenta-modeling platforms. Dev Cell 59, 776–792.e711 (2024). 10.1016/j.devcel.2024.01.023

6 Damsky, C. H. et al. Integrin switching regulates normal trophoblast invasion. Development 120, 3657–3666 (1994). 10.1242/dev.120.12.3657

7 Kabir-Salmani, M., Shiokawa, S., Akimoto, Y., Sakai, K. & Iwashita, M. The role of alpha(5)beta(1)-integrin in the IGF-I-induced migration of extravillous trophoblast cells during the process of implantation. Mol Hum Reprod 10, 91–97 (2004). 10.1093/molehr/gah014

8 Damsky, C. H., Fitzgerald, M. L. & Fisher, S. J. Distribution patterns of extracellular matrix components and adhesion receptors are intricately modulated during first trimester cytotrophoblast differentiation along the invasive pathway, in vivo. J Clin Invest 89, 210–222 (1992). 10.1172/jci115565

9 Ackerman Iv, W. E., et al. Epigenetic Changes Regulating Epithelial-Mesenchymal Plasticity in Human Trophoblast Differentiation. Cells 14 (2025). 10.3390/cells14130970

10 Cindrova-Davies, T. & Sferruzzi-Perri, A. N. Human placental development and function. Semin Cell Dev Biol 131, 66–77 (2022). 10.1016/j.semcdb.2022.03.039

11 Pollheimer, J. et al. Activation of the canonical wingless/T-cell factor signaling pathway promotes invasive differentiation of human trophoblast. Am J Pathol 168, 1134–1147 (2006). 10.2353/ajpath.2006.050686

12 Zhao, H. B. et al. E-cadherin, as a negative regulator of invasive behavior of human trophoblast cells, is down-regulated by cyclosporin A via epidermal growth factor/extracellular signal-regulated protein kinase signaling pathway. Biol Reprod 83, 370–376 (2010). 10.1095/biolreprod.110.083402

13 Arimoto-Ishida, E. et al. Up-regulation of alpha5-integrin by E-cadherin loss in hypoxia and its key role in the migration of extravillous trophoblast cells during early implantation. Endocrinology 150, 4306–4315 (2009). 10.1210/en.2008-1662

14 Bax, B. E. & Bloxam, D. L. Energy metabolism and glycolysis in human placental trophoblast cells during differentiation. Biochim Biophys Acta 1319, 283–292 (1997). 10.1016/s0005-2728(96)00169-7

15 Liu, X. et al. Down-regulation of PDK4 is Critical for the Switch of Carbohydrate Catabolism during Syncytialization of Human Placental Trophoblasts. Sci Rep 7, 8474 (2017). 10.1038/s41598-017-09163-8

16 Rodesch, F., Simon, P., Donner, C. & Jauniaux, E. Oxygen measurements in endometrial and trophoblastic tissues during early pregnancy. Obstet Gynecol 80, 283–285 (1992).

17 Jauniaux, E. et al. Onset of maternal arterial blood flow and placental oxidative stress. A possible factor in human early pregnancy failure. Am J Pathol 157, 2111–2122 (2000). 10.1016/s0002-9440(10)64849-3

18 Jauniaux, E., Watson, A. & Burton, G. Evaluation of respiratory gases and acid-base gradients in human fetal fluids and uteroplacental tissue between 7 and 16 weeks’ gestation. Am J Obstet Gynecol 184, 998–1003 (2001). 10.1067/mob.2001.111935

19 Withington, S. L. et al. Loss of Cited2 affects trophoblast formation and vascularization of the mouse placenta. Developmental biology 294, 67–82 (2006). 10.1016/j.ydbio.2006.02.025

20 Okazaki, K. & Maltepe, E. Oxygen, epigenetics and stem cell fate. Regen Med 1, 71–83 (2006). 10.2217/17460751.1.1.71

21 Adelman, D. M., Gertsenstein, M., Nagy, A., Simon, M. C. & Maltepe, E. Placental cell fates are regulated in vivo by HIF-mediated hypoxia responses. Genes & development 14, 3191–3203 (2000). 10.1101/gad.853700

22 Genbacev, O., Zhou, Y., Ludlow, J. W. & Fisher, S. J. Regulation of human placental development by oxygen tension. Science 277, 1669–1672 (1997). 10.1126/science.277.5332.1669

23 Wakeland, A. K. et al. Hypoxia Directs Human Extravillous Trophoblast Differentiation in a Hypoxia-Inducible Factor-Dependent Manner. Am J Pathol 187, 767–780 (2017). 10.1016/j.ajpath.2016.11.018

24 Jaremek, A. et al. Genome-Wide Analysis of Hypoxia-Inducible Factor Binding Reveals Targets Implicated in Impaired Human Placental Syncytiotrophoblast Formation under Low Oxygen. Am J Pathol 193, 846–865 (2023). 10.1016/j.ajpath.2023.03.006

25 Zhou, H. et al. Regulators involved in trophoblast syncytialization in the placenta of intrauterine growth restriction. Front Endocrinol (Lausanne) 14, 1107182 (2023). 10.3389/fendo.2023.1107182

26 Caniggia, I. et al. Hypoxia-inducible factor-1 mediates the biological effects of oxygen on human trophoblast differentiation through TGFbeta(3). J Clin Invest 105, 577–587 (2000). 10.1172/jci8316

27 Genbacev, O., Joslin, R., Damsky, C. H., Polliotti, B. M. & Fisher, S. J. Hypoxia alters early gestation human cytotrophoblast differentiation/invasion in vitro and models the placental defects that occur in preeclampsia. J Clin Invest 97, 540–550 (1996). 10.1172/jci118447

28 James, J. L., Stone, P. R. & Chamley, L. W. The effects of oxygen concentration and gestational age on extravillous trophoblast outgrowth in a human first trimester villous explant model. Hum Reprod 21, 2699–2705 (2006). 10.1093/humrep/del212

29 James, J. L., Stone, P. R. & Chamley, L. W. The regulation of trophoblast differentiation by oxygen in the first trimester of pregnancy. Hum Reprod Update 12, 137–144 (2006). 10.1093/humupd/dmi043

30 Lash, G. E. et al. Low oxygen concentrations inhibit trophoblast cell invasion from early gestation placental explants via alterations in levels of the urokinase plasminogen activator system. Biol Reprod 74, 403–409 (2006). 10.1095/biolreprod.105.047332

31 Treissman, J. et al. Low oxygen enhances trophoblast column growth by potentiating differentiation of the extravillous lineage and promoting LOX activity. Development 147 (2020). 10.1242/dev.181263

32 Chang, C. W., Wakeland, A. K. & Parast, M. M. Trophoblast lineage specification, differentiation and their regulation by oxygen tension. J Endocrinol 236, R43–r56 (2018). 10.1530/joe-17-0402

33 Goldman-Wohl, D. & Yagel, S. Regulation of trophoblast invasion: from normal implantation to pre-eclampsia. Mol Cell Endocrinol 187, 233–238 (2002). 10.1016/s0303-7207(01)00687-6

34 Wang, G. L., Jiang, B. H., Rue, E. A. & Semenza, G. L. Hypoxia-inducible factor 1 is a basic-helix-loop-helix-PAS heterodimer regulated by cellular O2 tension. Proc Natl Acad Sci U S A 92, 5510–5514 (1995). 10.1073/pnas.92.12.5510

35 Fryer, B. H. & Simon, M. C. Hypoxia, HIF and the placenta. Cell Cycle 5, 495–498 (2006). 10.4161/cc.5.5.2497

36 Dunwoodie, S. L. The role of hypoxia in development of the Mammalian embryo. Dev Cell 17, 755–773 (2009). 10.1016/j.devcel.2009.11.008

37 Hemberger, M., Hanna, C. W. & Dean, W. Mechanisms of early placental development in mouse and humans. Nature Reviews Genetics 21, 27–43 (2020). 10.1038/s41576-019-0169-4

38 Gerri, C. et al. Initiation of a conserved trophectoderm program in human, cow and mouse embryos. Nature 587, 443–447 (2020). 10.1038/s41586-020-2759-x

39 Gerri, C., Menchero, S., Mahadevaiah, S. K., Turner, J. M. A. & Niakan, K. K. Human Embryogenesis: A Comparative Perspective. Annu Rev Cell Dev Biol 36, 411–440 (2020). 10.1146/annurev-cellbio-022020-024900

40 Soncin, F., Natale, D. & Parast, M. M. Signaling pathways in mouse and human trophoblast differentiation: a comparative review. Cell Mol Life Sci 72, 1291–1302 (2015). 10.1007/s00018-014-1794-x

41 Okae, H. et al. Derivation of Human Trophoblast Stem Cells. Cell Stem Cell 22, 50–63.e56 (2018). 10.1016/j.stem.2017.11.004

42 Jokimaa, V. et al. Expression of syndecan-1 in human placenta and decidua. Placenta 19, 157–163 (1998). 10.1016/s0143-4004(98)90004-2

43 Yang, Y. et al. VGLL1 cooperates with TEAD4 to control human trophectoderm lineage specification. Nat Commun 15, 583 (2024). 10.1038/s41467-024-44780-8

44 Meinhardt, G. et al. Wnt-dependent T-cell factor-4 controls human etravillous trophoblast motility. Endocrinology 155, 1908–1920 (2014). 10.1210/en.2013-2042

45 Haider, S. et al. Self-Renewing Trophoblast Organoids Recapitulate the Developmental Program of the Early Human Placenta. Stem Cell Reports 11, 537–551 (2018). 10.1016/j.stemcr.2018.07.004

46 Turco, M. Y. et al. Trophoblast organoids as a model for maternal–fetal interactions during human placentation. Nature 564, 263–267 (2018). 10.1038/s41586-018-0753-3

47 Law, R. H. et al. An overview of the serpin superfamily. Genome Biol 7, 216 (2006). 10.1186/gb-2006-7-5-216

48 Ichikawa, M. K. et al. Ets family proteins regulate the EMT transcription factors Snail and ZEB in cancer cells. FEBS Open Bio 12, 1353–1364 (2022). 10.1002/2211-5463.13415

49 Schubert, M. et al. Perturbation-response genes reveal signaling footprints in cancer gene expression. Nat Commun 9, 20 (2018). 10.1038/s41467-017-02391-6

50 Apps, R. et al. Human leucocyte antigen (HLA) expression of primary trophoblast cells and placental cell lines, determined using single antigen beads to characterize allotype specificities of anti-HLA antibodies. Immunology 127, 26–39 (2009). 10.1111/j.1365-2567.2008.03019.x

51 Zhao, H. J. et al. Bone morphogenetic protein 2 promotes human trophoblast cell invasion and endothelial-like tube formation through ID1-mediated upregulation of IGF binding protein-3. Faseb j 34, 3151–3164 (2020). 10.1096/fj.201902168RR

52 Wang, Y., Li, Y. & Nie, G. HtrA4 is well conserved only in higher primates and functionally important for EVT differentiation. Placenta 152, 53–64 (2024). 10.1016/j.placenta.2024.05.132

53 DeWaal, D. et al. Hexokinase-2 depletion inhibits glycolysis and induces oxidative phosphorylation in hepatocellular carcinoma and sensitizes to metformin. Nat Commun 9, 446 (2018). 10.1038/s41467-017-02733-4

54 Griggs, L. A. et al. Fibronectin fibrils regulate TGF-β1-induced Epithelial-Mesenchymal Transition. Matrix Biol 60-61, 157–175 (2017). 10.1016/j.matbio.2017.01.001

55 Chesney, J. et al. Fructose-2,6-bisphosphate synthesis by 6-phosphofructo-2-kinase/fructose-2,6-bisphosphatase 4 (PFKFB4) is required for the glycolytic response to hypoxia and tumor growth. Oncotarget 5, 6670–6686 (2014). 10.18632/oncotarget.2213

56 Das, S., Das, S., Maity, A. & Maiti, S. Mechanistic Insights into FNBP4-Mediated Regulation of non-diaphanous formin FMN1 in Actin Cytoskeleton Dynamics. bioRxiv, 2024.2012.2007.627365 (2024). 10.1101/2024.12.07.627365

57 Palmer, N. J., Barrie, K. R. & Dominguez, R. Mechanisms of actin filament severing and elongation by formins. Nature 632, 437–442 (2024). 10.1038/s41586-024-07637-0

58 Chinthalapudi, K., Rangarajan, E. S. & Izard, T. The interaction of talin with the cell membrane is essential for integrin activation and focal adhesion formation. Proc Natl Acad Sci U S A 115, 10339–10344 (2018). 10.1073/pnas.1806275115

59 Caniggia, I., Winter, J., Lye, S. J. & Post, M. Oxygen and placental development during the first trimester: implications for the pathophysiology of pre-eclampsia. Placenta 21 **Suppl A**, S25–30 (2000). 10.1053/plac.1999.0522

60 Rajakumar, A., Brandon, H. M., Daftary, A., Ness, R. & Conrad, K. P. Evidence for the functional activity of hypoxia-inducible transcription factors overexpressed in preeclamptic placentae. Placenta 25, 763–769 (2004). 10.1016/j.placenta.2004.02.011

61 Zamudio, S. et al. Human placental hypoxia-inducible factor-1alpha expression correlates with clinical outcomes in chronic hypoxia in vivo. Am J Pathol 170, 2171–2179 (2007). 10.2353/ajpath.2007.061185

62 Zheng, X. et al. Repression of hypoxia-inducible factor-1 contributes to increased mitochondrial reactive oxygen species production in diabetes. Elife 11 (2022). 10.7554/eLife.70714

63 Mukherjee, I. et al. Oxidative stress-induced impairment of trophoblast function causes preeclampsia through the unfolded protein response pathway. Sci Rep 11, 18415 (2021). 10.1038/s41598-021-97799-y

64 Lamers, M. L., Almeida, M. E., Vicente-Manzanares, M., Horwitz, A. F. & Santos, M. F. High glucose-mediated oxidative stress impairs cell migration. PLoS One 6, e22865 (2011). 10.1371/journal.pone.0022865

65 Semenza, G. L. Targeting HIF-1 for cancer therapy. Nat Rev Cancer 3, 721–732 (2003). 10.1038/nrc1187

66 Sheridan, M. A. et al. Establishment and differentiation of long-term trophoblast organoid cultures from the human placenta. Nat Protoc 15, 3441–3463 (2020). 10.1038/s41596-020-0381-x

67 Dekkers, J. F. et al. High-resolution 3D imaging of fixed and cleared organoids. Nat Protoc 14, 1756–1771 (2019). 10.1038/s41596-019-0160-8

68 Stringer, C., Wang, T., Michaelos, M. & Pachitariu, M. Cellpose: a generalist algorithm for cellular segmentation. Nat Methods 18, 100–106 (2021). 10.1038/s41592-020-01018-x

69 Zheng, G. X. et al. Massively parallel digital transcriptional profiling of single cells. Nat Commun 8, 14049 (2017). 10.1038/ncomms14049

70 Hao, Y. et al. Integrated analysis of multimodal single-cell data. Cell 184, 3573–3587.e3529 (2021). 10.1016/j.cell.2021.04.048

71 Hafemeister, C. & Satija, R. Normalization and variance stabilization of single-cell RNA-seq data using regularized negative binomial regression. Genome Biol 20, 296 (2019). 10.1186/s13059-019-1874-1

72 Marsh, S. E. et al. Dissection of artifactual and confounding glial signatures by single-cell sequencing of mouse and human brain. Nat Neurosci 25, 306–316 (2022). 10.1038/s41593-022-01022-8

73 van den Brink, S. C., et al. Single-cell sequencing reveals dissociation-induced gene expression in tissue subpopulations. Nat Methods 14, 935–936 (2017). 10.1038/nmeth.4437

74 Vento-Tormo, R. et al. Single-cell reconstruction of the early maternal–fetal interface in humans. Nature 563, 347–353 (2018). 10.1038/s41586-018-0698-6

